# DUX4-induced bidirectional HSATII satellite repeat transcripts form intranuclear double stranded RNA foci in human cell models of FSHD

**DOI:** 10.1101/734541

**Authors:** Sean C. Shadle, Sean R. Bennett, Chao-Jen Wong, Nancy A. Karreman, Amy E. Campbell, Silvère M. van der Maarel, Brenda L. Bass, Stephen J. Tapscott

## Abstract

The DUX4 transcription factor is normally expressed in the cleavage stage embryo and regulates genes involved in embryonic genome activation. Mis-expression of DUX4 in skeletal muscle, however, is toxic and causes facioscapulohumeral muscular dystrophy (FSHD). We recently showed DUX4-induced toxicity is due, in part, to the activation of the double-stranded RNA (dsRNA) response pathway and the accumulation of intranuclear dsRNA foci. Here, we determined the composition of DUX4-induced dsRNAs. We found that a subset of DUX4-induced dsRNAs originate from inverted *Alu* repeats embedded within the introns of DUX4-induced transcripts and from DUX4-induced dsRNA-forming intergenic transcripts enriched for endogenous retroviruses, *Alu* and LINE-1 elements. However, these repeat classes were also represented in dsRNAs from cells not expressing DUX4. In contrast, pericentric human satellite II (HSATII) repeats formed a class of dsRNA specific to the DUX4 expressing cells. Further investigation revealed that DUX4 can initiate the bidirectional transcription of normally heterochromatin-silenced HSATII repeats. DUX4 induced HSATII RNAs co-localized with DUX4-induced nuclear dsRNA foci and with intranuclear aggregation of EIF4A3 and ADAR1. Finally, gapmer-mediated knockdown of HSATII transcripts depleted DUX4-induced intranuclear ribonucleoprotein aggregates and decreased DUX4-induced cell death, suggesting that HSATII formed dsRNAs contribute to DUX4 toxicity.

## Introduction

The Double Homeobox 4 (DUX4) transcription factor is normally expressed in the testis, likely the germline cells (1), and in the cleavage stage embryo coincident with embryonic genome activation (EGA) (2–5). In contrast, the mis-expression of DUX4 in skeletal muscle causes FSHD (1, 6, 7). In both embryonic stem cells and in skeletal muscle cells, expression of DUX4 activates the transcription of hundreds of genes that are characteristically expressed in the cleavage stage embryo. DUX4 also activates the expression of normally silenced repetitive elements that are transcribed during EGA such as endogenous retroviruses (ERVs) and the pericentric human satellite II (HSATII) repeats (3, 4, 8, 9). The expression of DUX4 in skeletal muscle cells is toxic and causes apoptosis (10–13), however, the specific molecular pathways leading to cell toxicity are not fully understood. Previous studies have suggested multiple, non-mutually exclusive mechanisms for DUX4 toxicity including an increased sensitivity to oxidative stress (12, 14), interference with PAX3/PAX7 (12, 15) and the formation of insoluble TDP-43 nuclear aggregates due to impaired protein turnover (16).

More recently, we reported that the expression of DUX4 in skeletal muscle cells resulted in the accumulation of intranuclear foci of dsRNAs (13). The accumulation of DUX4-induced dsRNAs correlated with PKR and eIF2α phosphorylation, both proapoptotic characteristics of the cellular innate immune response typically triggered by viral dsRNAs (17). Indeed, knockdown of *EIF2AK2* (PKR) or *RNASEL*, another mediator of the dsRNA response pathway, decreased the amount of cell death in cells overexpressing DUX4 (13). However, the composition of the DUX4-induced intranuclear dsRNA foci was unknown.

In this study, we identified the composition of the dsRNAs induced following DUX4 expression in human skeletal muscle cells. In contrast to the mainly intronic origin of endogenous human dsRNAs (18–20), we found that a large portion of DUX4-induced dsRNAs originate from intergenic, non-protein coding regions of the genome, and were highly enriched for HSATII satellite sequences. We found that DUX4 initially induces the transcription of one strand of HSATII, which forms single-stranded intranuclear foci that co-localize with EIF4A3. Subsequently, the induction of the complementary HSATII strand results in dsRNA foci that co-localize with the dsRNA-binding ADAR1 enzyme. Finally, gapmer-mediated depletion of HSATII transcripts results in the loss of dsRNA foci and corresponding ribonucleoprotein aggregates, diminishes PKR phosphorylation, and decreases DUX4-induced cell death, suggesting that HSATII transcription contributes to DUX4 toxicity. Moreover, our results provide the first comprehensive study of double-stranded RNAs in non-affected and FSHD-affected human muscle cells.

## Results

### Double-stranded RNA immunoprecipitation and sequencing identifies regions of dsRNA induced by DUX4

Previous studies have identified double-stranded RNAs by high-throughput sequencing of immunoprecipitated RNAs (dsRIP-seq) that were pulled down with the widely used J2 dsRNA antibody (19, 21, 22). To identify dsRNAs induced by DUX4, we performed dsRIP-seq on a doxycycline-inducible DUX4 human myoblast cell line (MB135-iDUX4) with both the J2 and the independent monoclonal dsRNA-recognizing antibody, K1. After read mapping, we used the MACS2 peak-calling algorithm to compare the enrichment of the J2 or K1 dsRNA IPs to the mock IgG immunoprecipitation in order to identify dsRNA-enriched regions within each -/+ doxycycline (DOX) condition (Figure 1A). A correlation heatmap of read count data showed high similarity between the K1 and J2 antibodies within each condition (Supplementary Material, Figure S1), indicating that both antibodies identified a largely overlapping set of dsRNAs. Therefore, we combined the J2 and K1 data to identify candidate dsRNA regions for subsequent analyses.

**Figure 1.**
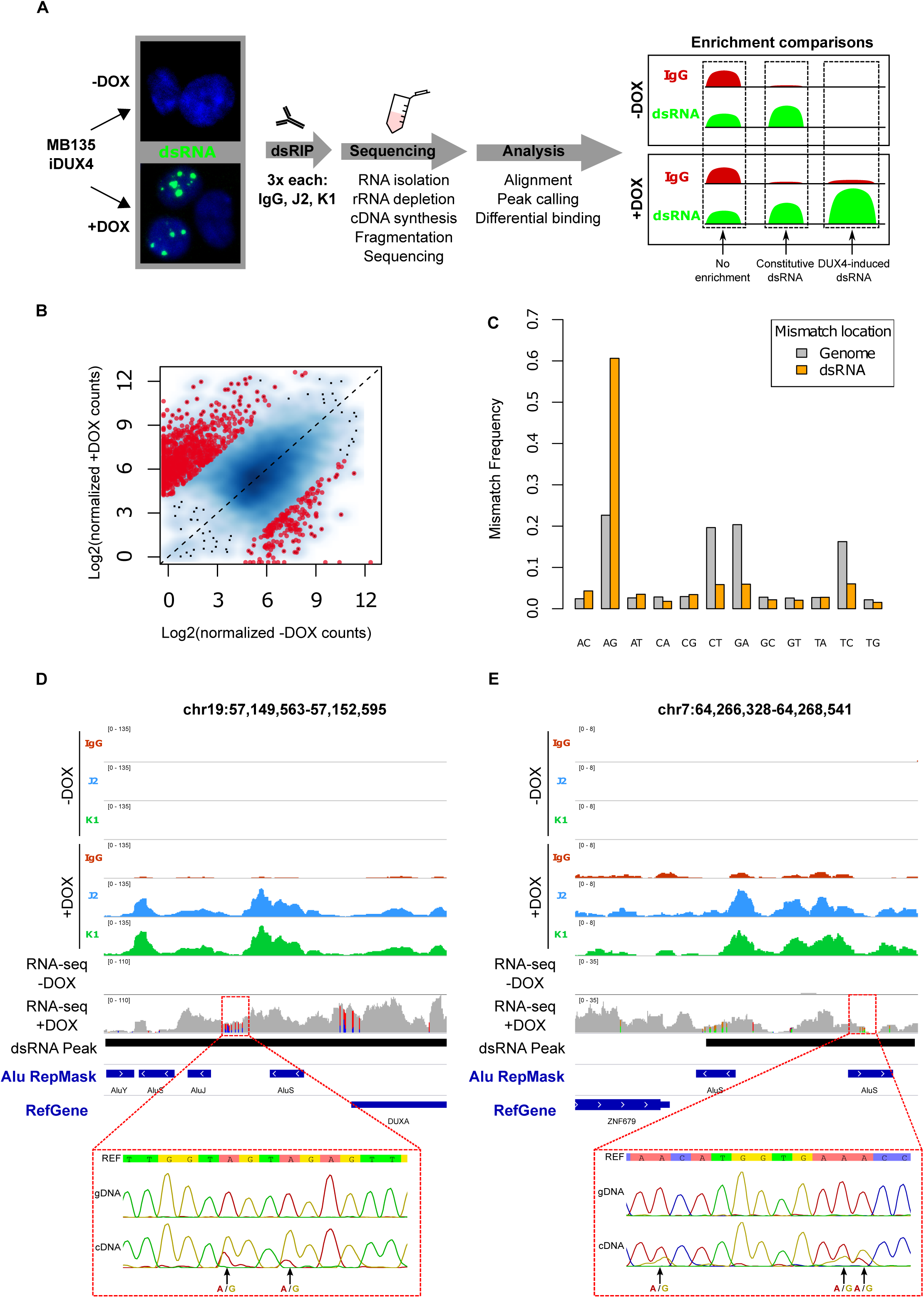
Double-stranded RNA immunoprecipitation and sequencing of DUX4-induced dsRNAs enriches for A-I edited transcripts. (**A**) Schematic of experimental outline for dsRNA immunoprecipitation and sequencing. Briefly, -/+ doxycycline (DOX) induced MB135-iDUX4 cells were lysed and subjected to immunoprecipitation of either matched antibody isotype control (IgG) or J2 or K1 antibodies, performed in triplicate. Libraries were constructed from isolated RNA and informatics analysis consisted of broad peak calling followed by differential enrichment testing. Within each (-/+ DOX) condition, dsRNA “enrichment” was determined against the IgG control, which essentially represents total RNA abundance. Comparison of enrichment between conditions provides a way to measure level of induction following DUX4 expression. Three separate cultures of MB135-iDUX4 cells were used per condition. (**B**) Scatterplot comparing normalized levels of dsRNAs in -/+ DOX conditions, depicted as the log2 counts plus a pseudocount value of 1.0. Red points highlight dsRNAs that show evidence for statistically significant differential enrichment and are limited to values with a threshold absolute log2 fold change of 4.0 and FDR-adjusted p-value of 1×10^-10^. (**C**) Mismatch frequencies of each of the 12 possible mismatch types within DUX4-induced dsRNAs in the +DOX stranded RNA-seq dataset within the indicated locations. (**D-E**) Browser screenshots in the indicated hg38 regions of normalized read counts for -/+ DOX immunoprecipitations. These tracks represent merged read counts for the indicated triplicate datasets. “RNA-seq -/+ DOX” refers to the stranded RNA-seq dataset and is shown as raw read counts such that mismatches (highlighted in non-grey colors) can be visualized. DUX4-induced dsRNA regions are depicted by black bars. Sanger sequencing results of amplicons designed across the annotated regions (red dashed lines) in either genomic DNA (gDNA) or complementary DNA (cDNA) of +DOX MB135-iDUX4 cells, with AG mismatches highlighted. Here, the reference strand is shown as the strand of origin for the transcripts, based on the stranded RNA-seq dataset.

To describe the genomic regions producing dsRNAs induced by DUX4, we compared transcripts differentially enriched between-DOX and +DOX conditions. Using an absolute log2 fold-change threshold of 4.0 and an FDR-adjusted p-value of 10^-10^, 870 regions showed increased dsRNA enrichment in DUX4 expressing cells (see (Supplementary Material Figure S2 for chromosomal locations) compared to 193 regions which showed decreased enrichment (Figure 1B and Supplementary Material Table S1), confirming a general induction of dsRNA enriched regions following DUX4 expression.

To verify that we successfully enriched for dsRNAs, we next examined dsRNA-specific RNA editing. The ADAR enzymes convert adenosine to inosine specifically in dsRNAs which manifests as A to G mismatching in RNA-seq reads (23). We therefore examined mismatches in a strand specific RNA-seq dataset of MB135-iDUX4 myoblasts expressing DUX4. Within DUX4-induced dsRNA regions, the A-G mismatch type accounted for more than 60% of all 12 possible called mismatches compared to the ∼22% frequency of A-G mismatches called in the entire genome-mapped reads (Figure 1C), confirming that the K1 and J2 antibodies successfully enriched for dsRNAs. We further confirmed that the A-G mismatches represented editing events rather than rare polymorphisms by topo-cloning and comparing the DNA sequence of MB135-iDUX4 cells with the RNA sequence in selected dsRNA regions (Figures 1D-1E and Supplementary Material Figure S3). Combined, the above results validate our dsRNA IP strategy and indicate, as previously observed (13), that DUX4 expression leads to the induction of dsRNA transcripts that are subject to dsRNA-specific ADAR editing.

### DUX4-induced double-stranded RNAs are enriched for non-coding intergenic RNAs

We next determined the annotated genomic features associated with regions of DUX4-induced dsRNAs. Constitutive dsRNAs, which were expressed in the presence or absence of DUX4 expression (Figure 2A), recapitulated the known intronic enrichment as well as slight 3-prime UTR enrichment for A-I edited dsRNA (Figure 2B and references (18–20)). Conversely, DUX4-induced dsRNAs had a profile shifted largely towards intergenic regions (Figure 2B), either upstream or downstream of the annotated gene, such as in the example shown upstream of *TP53BP2* (Figure 2C) or previous examples downstream of *DUXA* (Figure 1D) and *ZNF679* (Figure 1E), or embedded within larger DUX4-induced long non-coding intergenic transcripts that extended upwards of 500 kb in length (Figure 2D and Supplementary Material Figure S4).

**Figure 2.**
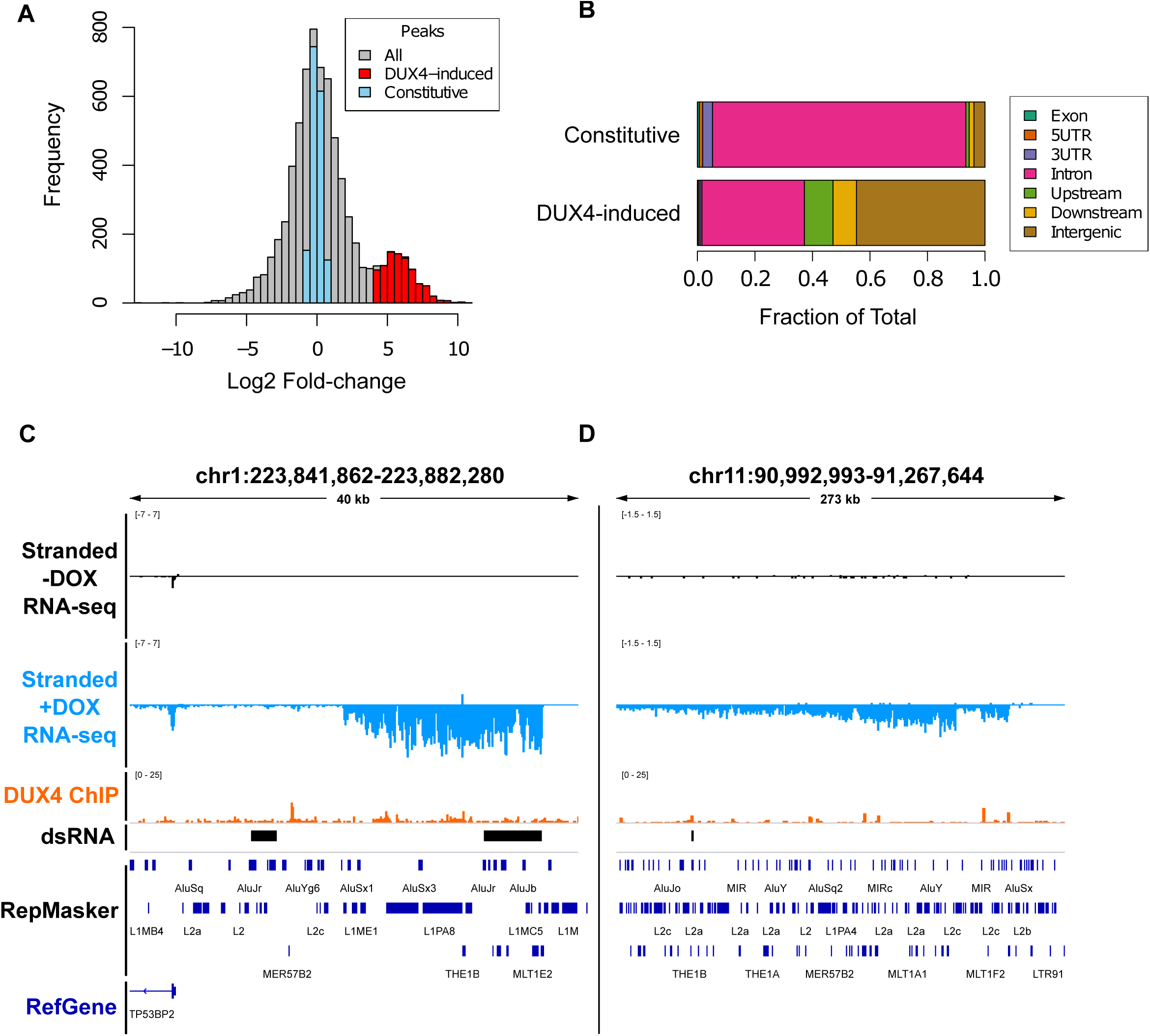
DUX4-induced double-stranded RNAs are enriched for non-coding intergenic RNAs. (**A**) Histogram of consensus “peaks” from DiffBind analysis of dsRIP-seq experiment with constitutive peaks (blue, |log2 fold change| < 1.0 and FDR-adjusted p-value > 0.05), and DUX4-increased (red, log2 fold change > 4.0 and FDR-adjusted p-value ≤ 1×10^-10^). (**B**) Genomic feature distribution overlap of each dsRNA, defined as in Figure 2A. See Methods for feature definitions and prioritizations. (**C-D**) Browser screenshots in the indicated hg38 regions of normalized stranded ribominus MB135-iDUX4 RNA-seq data in two intergenic locations. These tracks represent merged read counts for the indicated duplicate datasets.

Based on previously published ChIP-seq (8), many of these transcripts showed evidence for binding by DUX4 at endogenous retroviral LTR elements near the beginning of the transcript. Indeed, DUX4 bound significantly closer (p = 6.83×10^-10^) to DUX4-induced dsRNA regions than constitutive dsRNA regions (Supplementary Material Figure S5), suggesting a direct activation for many of the identified DUX4-induced dsRNAs.

Collectively, these data indicate that, in contrast to the largely intronic origin of dsRNAs in mammalian cells, a substantial portion of DUX4-induced dsRNAs are comprised of sequences embedded within non-coding, likely heterochromatin-enriched intergenic DUX4-induced transcripts. At least some of these transcripts appear to be directly activated by DUX4 as they originate in close proximity to DUX4 binding sites.

### DUX4-induced dsRNAs are enriched for repeat sequences including *Alu*, LINE-1, HERVL/MaLR and the pericentric HSATII repeat

Transcripts that contain repetitive sequences are known to form dsRNAs. For example, *Alu* SINE repeats are vastly overrepresented in human dsRNAs (18) where they often form inverted, hybridized pairs within a single RNA transcript (20, 24). LTR-containing ERVs have also been shown to form dsRNAs in tumor cells treated with 5-azacytidine (25, 26). To determine the set of repeats activated by DUX4, we used the Dfam database of repetitive elements (27), which identified the classes of repeats induced in our stranded total RNA sequencing dataset (Figure 3A and Supplementary Material Table S2). This analysis confirmed the previously published observation that DUX4 activates expression of HSATII pericentric satellite repeats, MaLR sequences such as THE1D, HERVL and a subset of LINE-1 repeats (3, 4, 8, 9, 28).

**Figure 3.**
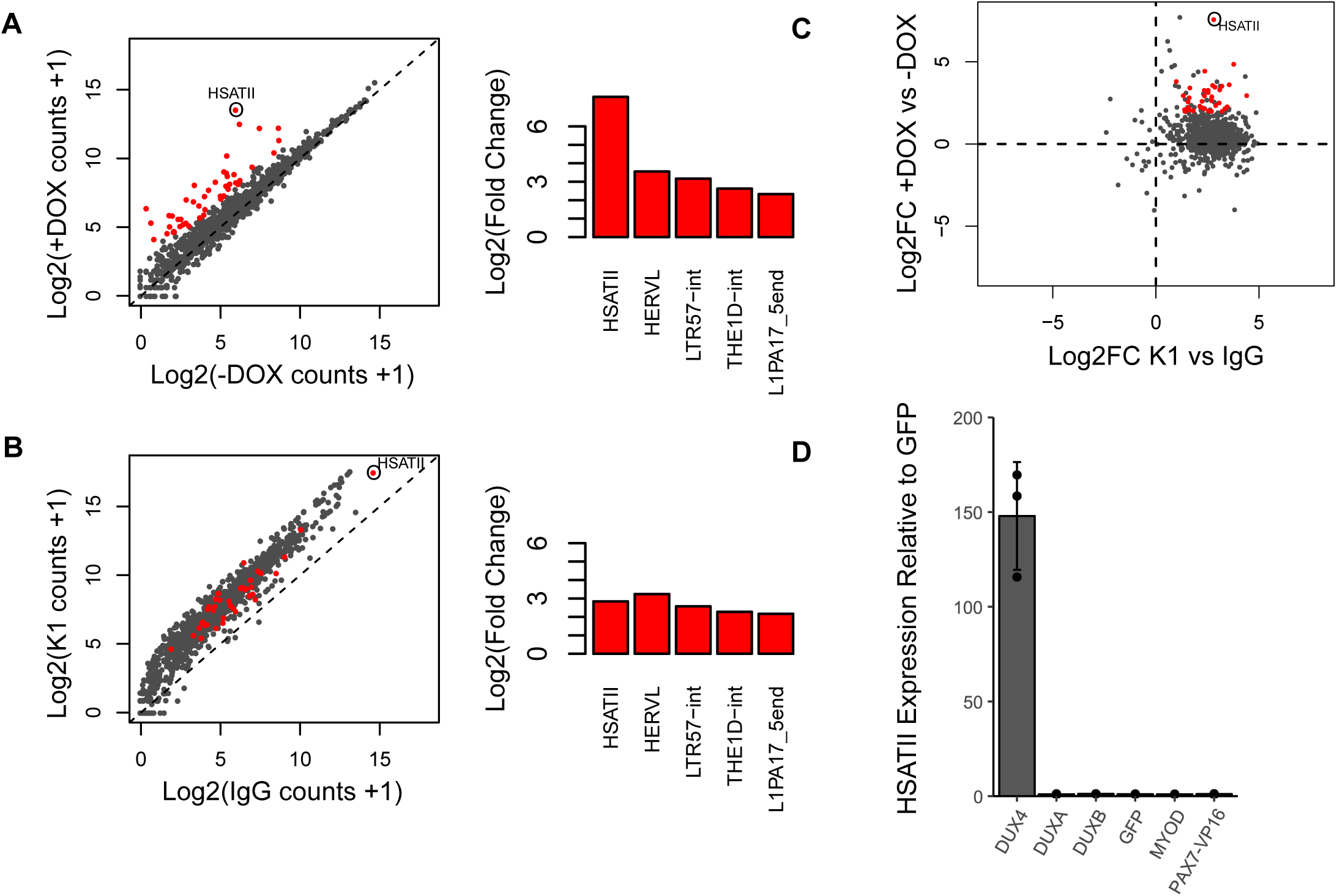
DUX4-induced dsRNAs are enriched for repeat sequences including HSATII. (**A**) Scatterplot depicting log2 normalized read counts of Dfam predicted repeat class subfamilies within stranded RNA-seq dataset of cells -/+ DOX is shown on the left. Repeat subfamilies are highlighted as red when |log2 fold change| > 2.0 and FDR-adjusted p-value < 0.01. DESeq2 moderated log2 fold change values of select DUX4-induced repeat subfamilies are shown on the right. (**B**) Scatterplot depicting log2 normalized read counts of Dfam predicted repeat class subfamilies within K1 vs IgG dsRIP-seq datasets in the +DOX condition. Significantly differentially expressed repeats (red dots) from (A) are considered enriched and highlighted as red in this plot if they meet the criteria of |log2 fold change| > 1.0 and FDR-adjusted p-value < 0.01. DESeq2 moderated log2 fold change values of select repeat subfamilies are shown on the right. (**C**) Scatterplot of Dfam predicted repeat class subfamilies showing DESeq2 moderated log2 fold change values (Log2FC) in +DOX vs-DOX stranded RNA-seq (y-axis) compared to moderated log2 fold change values in K1 vs IgG RNA immunoprecipitations in the +DOX condition (x-axis). Red points are highlighted as in panel (B). (**D**) RT-qPCR showing levels of endogenous HSATII RNA expression relative to GFP following transfection of HEK293T cells with the indicated expression vectors (X-axis). All indicated expression vectors were cloned into the pCS2 backbone. Error bars represent standard deviation of the mean of three separate cultures, depicted as individual points.

We next determined repeat subfamilies that were enriched by the K1 or J2 antibodies compared to the IgG control in doxycycline treated MB135-iDUX4i cells (Supplementary Material Table S3). Intersecting all dsRNA enriched repeats with DUX4-induced repeats identified the set of repeats that were both strongly induced by DUX4 and were also enriched for dsRNAs (red dots, Figures 3B-3C and Supplementary Material Figure S6A-S6B). Validating this analysis, *Alu* repeat subfamilies were enriched in the dsRIP-seq datasets compared to the IgG control, though, as expected, they were not specifically upregulated following DUX4 expression (Supplementary Material Figure S7A). LINE-1, ERVL-MaLR and HERVL repeats were also modestly enriched in the K1 and J2 RIPs (Supplementary Material Figures S7B-S7C and Supplementary Material Table S3) and showed increased expression by DUX4 (Supplementary Material Table S2). However, the most highly induced repeat by DUX4 which was also enriched for dsRNA was the HSATII class of pericentric satellite repeats (Figures 3A-3C and Supplementary Material Figures S6A-S6B).

Compared to non-DUX4 expressing cells where 0.00-0.01% of dsRIP-seq reads mapped to HSATII, approximately 4.1% of K1 dsRIP-seq reads from DUX4 expressing myoblasts aligned to HSATII repeats. For unknown reasons, the J2 antibody showed more modest affinity than K1 for HSATII (compare Figure 3B and Supplementary Material Figure S6A), though these two antibodies do have differences in dsRNA epitope preferences (29). Intriguingly, HSATII repeat transcripts are known to form intranuclear foci after aberrant de-repression in certain cancer cells, appearing as distinct spots (30), reminiscent of our DUX4-induced dsRNA foci. For these reasons, we next focused on determining whether DUX4-induced intranuclear dsRNA foci were comprised of HSATII transcripts.

### DUX4 binds to and activates HSATII transcription

Satellite repeats can be transcribed during certain cellular stresses such as heat shock (31). However, our previously published ChIP-seq indicated that DUX4 directly binds HSATII repeats (3, 9). The consensus HSATII sequence contains close matches to the known DUX4 binding motif (Supplementary Material Figure S8A) and variation within unmapped HSATII repeat sequences (9, 32) creates the potential for large regions of arrayed near-perfect matches for DUX4 binding sites within pericentric regions. Indeed, Sanger sequencing of cloned HSATII PCR amplicons verified that these repeats often contain multiple copies of the consensus DUX4 binding motif (Supplementary Material Figure S8A), suggesting that DUX4 binding to the repeats might directly induce HSATII transcription, leading to dsRNA formation.

To more directly test whether DUX4 can activate HSATII transcription, we cloned a multicopy HSATII repeat that contained DUX4 binding motifs into the promoterless pGL3 luciferase reporter vector in both the forward and reverse orientation, relative to the consensus HSATII sequence (Supplementary Material Figure S8B). Co-transfection of HEK293T cells with this HSATII-luciferase reporter and a DUX4 expression vector or control GFP expression vector showed robust activation of luciferase RNA expression by DUX4 which was dependent on the presence of the HSATII sequence, but largely independent of the orientation (Supplementary Material Figure S8C). Further demonstrating the specificity of HSATII activation by DUX4, transcriptional upregulation of endogenous HSATII was observed following over-expression of DUX4 but was not observed after over-expression of the related double homeobox proteins, DUXA or DUXB or other transcription factors MYOD or PAX7 fused to the VP16 (PAX7-VP16) transactivation domain (Figure 3D). These results provide additional evidence for the specificity of the binding and transcriptional activation of HSATII repeats by DUX4.

### Temporally controlled bidirectional HSATII transcription forms DUX4-induced nuclear dsRNA foci

Analysis of the stranded DUX4-induced RNA-seq data demonstrated that, although reads predominantly mapped to the reverse complement of the consensus HSATII sequence (hereafter referred to as “reverse” HSATII transcripts), a minor proportion of “forward” transcripts were also evident (Figure 4A). Moreover, HSATII repeats can be bidirectionally transcribed in the context of some tumor cells (33). For these reasons, we postulated that bidirectional HSATII transcription might lead to the accumulation of intermolecularly formed HSATII dsRNA foci in DUX4 expressing cells, and that transcription of the forward strand might be the rate-limiting step for dsRNA formation.

**Figure 4.**
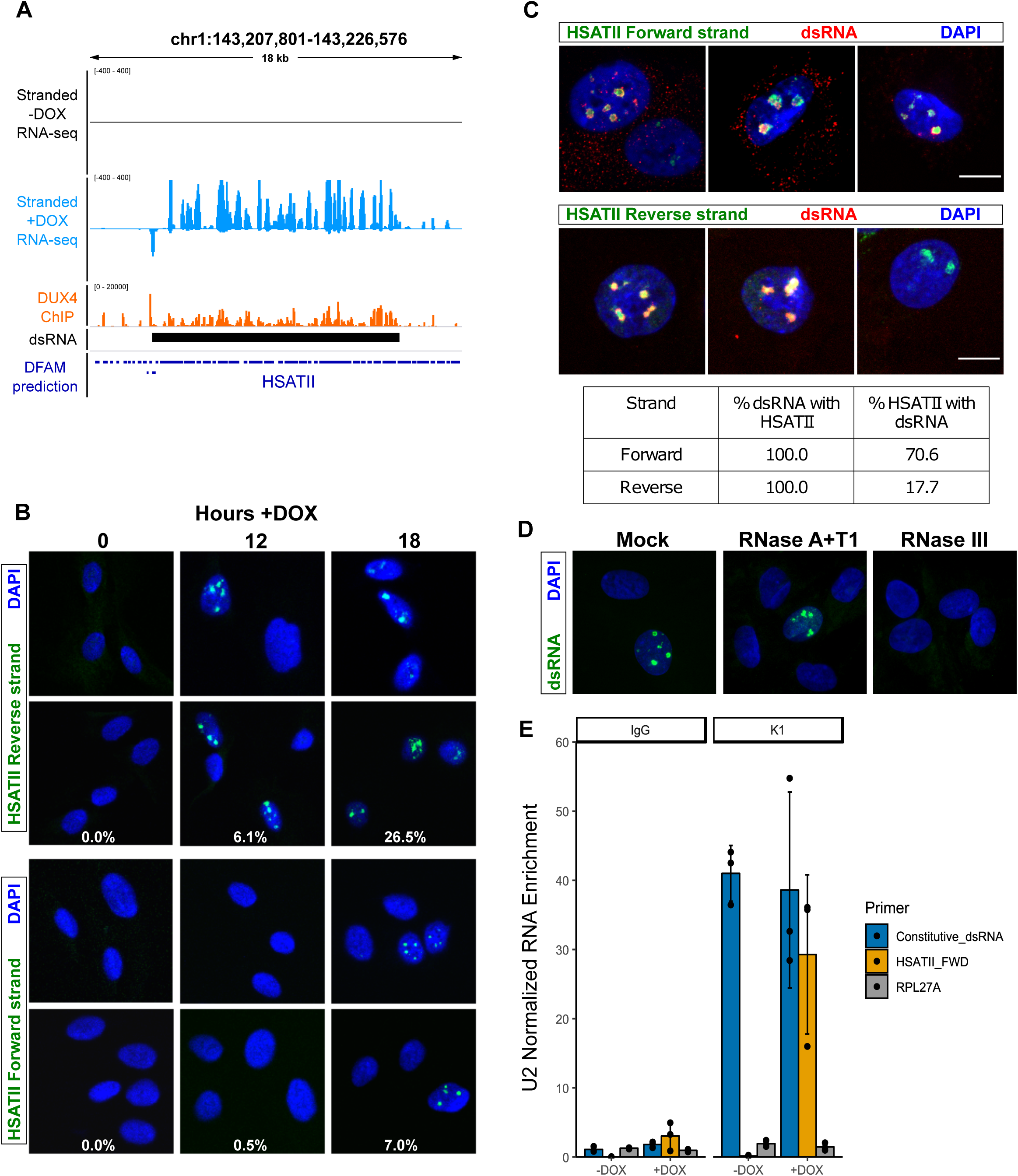
DUX4-induced HSATII transcripts are bidirectionally transcribed and form dsRNA. (**A**) Browser screenshot in the indicated hg38 region of normalized stranded ribominus MB135-iDUX4 RNA-seq data. Forward and reverse read counts are indicated above and below the horizontal line, respectively. Shown at the bottom are Dfam predicted HSATII repeat locations. (**B**) Confocal images of RNA fluorescence in situ hybridization of MB135-iDUX4 cells treated with doxycycline for the indicated times and hybridized with probes that detect the indicated HSATII strands, where forward and reverse is relative to the consensus sequence. Images are representative from two independent time-course experiments conducted on separate days. Estimates for percent of foci positive nuclei are indicated for each time point and are from > 150 random nuclei counted per time point. (**C**) Combined immunofluorescence in MB135-iDUX4 cells at 18 hours +DOX, using the K1 dsRNA antibody and RNA-FISH with probe detecting forward (upper) or reverse (lower) HSATII transcripts. Images are representative from two independent, combined IF RNA-FISH experiments conducted on separate days. Note, with the reverse probe a larger subset of DUX4 expressing cells contained HSATII foci, but no dsRNA foci (top), whereas another class contained HSATII foci which co-localized with dsRNA foci (bottom). Scale bars indicate 10 µm. Estimates for percentages of nuclear foci co-occurrence of the indicated HSATII strand and K1 antibody is shown in table below and were made from counts of > 250 randomly selected nuclei per time point. (**D**) K1 immunofluorescence in MB135-iDUX4 cells at 18 hours +DOX treated with the indicated RNase enzyme or mock treated prior to the immunofluorescence. No dsRNA foci were detected after treatment with RNase III, a double-stranded RNase. Experiment was performed twice, and a representative image is shown. (**E)** RT-qPCR of RNAs immunoprecipitations using K1 or IgG as control in MB135-iDUX4 cells -/+ DOX for approximately 18 hours, as indicated. Primers were designed to detect a constitutive dsRNA from our dsRIP-seq dataset or forward HSATII after strand-specific RT-qPCR. To account for differences in total RNA levels and reverse transcription efficiency, IPs were normalized to the amount of U2 RNA, which serves as an abundant background RNA species that should not be enriched by K1 levels. Error bars represent the standard deviation of the mean for three independent immunoprecipitations, which are shown as individual data points.

To determine the relationship of forward and reverse HSATII transcripts to the intranuclear dsRNA foci, we performed RNA fluorescence *in situ* hybridization (RNA-FISH) with probes detecting the forward or reverse HSATII transcripts (30) in a time course following doxycycline induction of MB135-iDUX4 cells. The probes to either the forward or reverse HSATII transcripts revealed intranuclear foci at the 18-hour time point following doxycycline treatment (Figure 4B), indicating that DUX4 induces bidirectional expression of HSATII. Interestingly, the probe targeting the reverse HSATII transcript identified foci in a substantial percentage of nuclei at both 12 and 18 hours following DUX4-induction, whereas the probe to the forward transcript identified obvious foci only at the 18-hour time point and in a smaller proportion of nuclei (Figure 4B).

Next, we performed immunofluorescence using the K1 antibody combined with HSATII RNA-FISH to determine whether these HSATII transcripts might overlap dsRNA foci. This experiment revealed that the DUX4-induced nuclear dsRNA foci nearly perfectly coincided with HSATII RNA foci (Figure 4C). These K1 stained foci were confirmed as double stranded because they were not evident following treatment with RNase III, a double-stranded ribonuclease, but were still evident following treatment with the single stranded RNases A and T1 (Figure 4D).

Because reverse strand HSATII foci formation were relatively more common and preceded forward strand HSATII foci formation, we predicted that forward transcripts would be more highly associated with dsRNA foci. Indeed, in a majority (∼70.6%) of nuclei, the forward HSATII foci coincided with dsRNA, whereas the reverse HSATII foci were more frequent and coincided with K1-positive dsRNA aggregates in a minority of nuclei (∼17.7%) (Figure 4C). Because all K1 dsRNA foci showed positive FISH signal for both forward and reverse HSATII RNA strands, our data indicate that DUX4-induced nuclear dsRNA aggregates were formed via bidirectional transcription of HSATII repeats, with the less abundant forward HSATII strand acting as a limiting factor for dsRNA formation.

To independently confirm that transcription of the forward HSATII strand is the rate-limiting step in the formation of DUX4-induced dsRNA foci, we performed strand-specific RT-qPCR following K1 dsRIP. This experiment revealed that forward HSATII transcripts were highly enriched by K1 compared to IgG in DUX4 expressing cells (Figure 4E), whereas the enrichment of HSATII in a non-strand-specific RT-qPCR was more modest (Supplementary Material Figure S9). This result is expected if forward HSATII transcripts are limiting for the formation of dsRNA. Combined, our results strongly suggest that DUX4-induced intranuclear dsRNA foci are comprised of temporally controlled bidirectional HSATII repeat transcripts that form upon the protracted expression of the forward HSATII strand.

### HSATII transcription is associated with EIF4A3 and ADAR1 aggregation and correlates with the formation of intranuclear BMI1 foci and phosphorylation of H2AX

We next sought to determine whether nuclear proteins may co-aggregate as a consequence of HSATII-containing dsRNA foci formation in DUX4 expressing cells. We previously showed that the exon junction complex factor EIF4A3 accumulates in intranuclear foci following DUX4 induction (13). In this prior study, EIF4A3 aggregates appeared earlier than dsRNA foci, but nearly all dsRNA foci co-localized with EIF4A3 foci upon their appearance. We therefore performed combined immunofluorescence and RNA-FISH of DUX4-induced cells with an antibody to EIF4A3 and the FISH probe to the HSATII reverse strand. This showed strong co-localization of EIF4A3 and reverse HSATII foci (Supplementary Material Figure S10A), indicating that EIF4A3 accumulates with reverse HSATII transcripts prior to the appearance of forward HSATII transcripts and the formation of dsRNA.

Although we cannot directly assess ADAR editing of HSATII repeats due to polymorphic variations from the consensus sequence (9), FISH combined with an antibody to ADAR1 indicated a strong redistribution of ADAR1 (which specifically binds dsRNA) to HSATII foci in DUX4 expressing cells (Supplementary Material Figure S10B). Nearly 100% of these ADAR1 foci also co-localized with K1 dsRNA nuclear staining (Supplementary Material Figure S10C), suggesting that DUX4-induced HSATII dsRNA is bound by ADAR1 and leads to ADAR1 accumulation as intranuclear foci. Cells with either EIF4A3 or ADAR1 foci had lost their normal pan-nuclear distribution, indicating a possible sequestration of these proteins in the foci.

We also wondered about other consequences of HSATII dsRNA formation in DUX4 expressing cells. Derepression of HSATII repeats in cancer cells has been associated with the formation of polycomb bodies (30, 34) and with double-strand DNA breaks (35). In our model system, DUX4 induction of HSATII repeats was similarly associated with the formation of polycomb bodies and foci of DNA damage, as determined by immunodetection of the PRC1 complex protein BMI1 and foci of phospho-H2AX (Supplementary Material Figure S11A). The BMI1 foci did not co-localize with induced HSATII RNA foci (Supplementary Material Figure S11B), but did show strong co-localization with the phospho-H2AX foci suggesting that the PRC1 complex was accumulating at sites of DNA damage, although it is possible that they might also be associated with demethylated HSATII DNA regions in the genome as has been suggested (30).

### HSATII transcripts are necessary for DUX4-induced dsRNA foci and contribute to toxicity

Locked nucleic acid gapmer oligonucleotides can degrade cellular transcripts via RNase H-dependent cleavage of complementary transcripts following hybridization, allowing for effective knockdown of nuclear transcripts, including satellite repeat RNAs (36). To determine whether DUX4-induced HSATII transcripts are necessary for dsRNA formation, we transfected MB135-iDUX4 cells with two separate pooled HSATII-targeted gapmers designed against both the forward and reverse HSATII RNA strands prior to doxycycline induction of DUX4. Our results demonstrated that transfection of HSATII-gapmers can deplete DUX4-induced HSATII transcripts as compared to a non-targeting control gapmer (Figure 5A), with no discernable effect on DUX4 expression (Supplementary Material Figure S12) or DUX4 target gene expression (Figure 5B). The F1R2 gapmer pool was more effective at depleting HSATII transcripts than the F2R1 pool (Figure 5A). Importantly, gapmer-mediated HSATII knockdown diminished the formation of dsRNA and EIF4A3 foci following DUX4 expression (Figure 5C), particularly in the case of the more effective F1R2 gapmer pool. These results indicate that HSATII expression is necessary for the formation of DUX4-induced intranuclear dsRNA and EIF4A3 foci.

**Figure 5.**
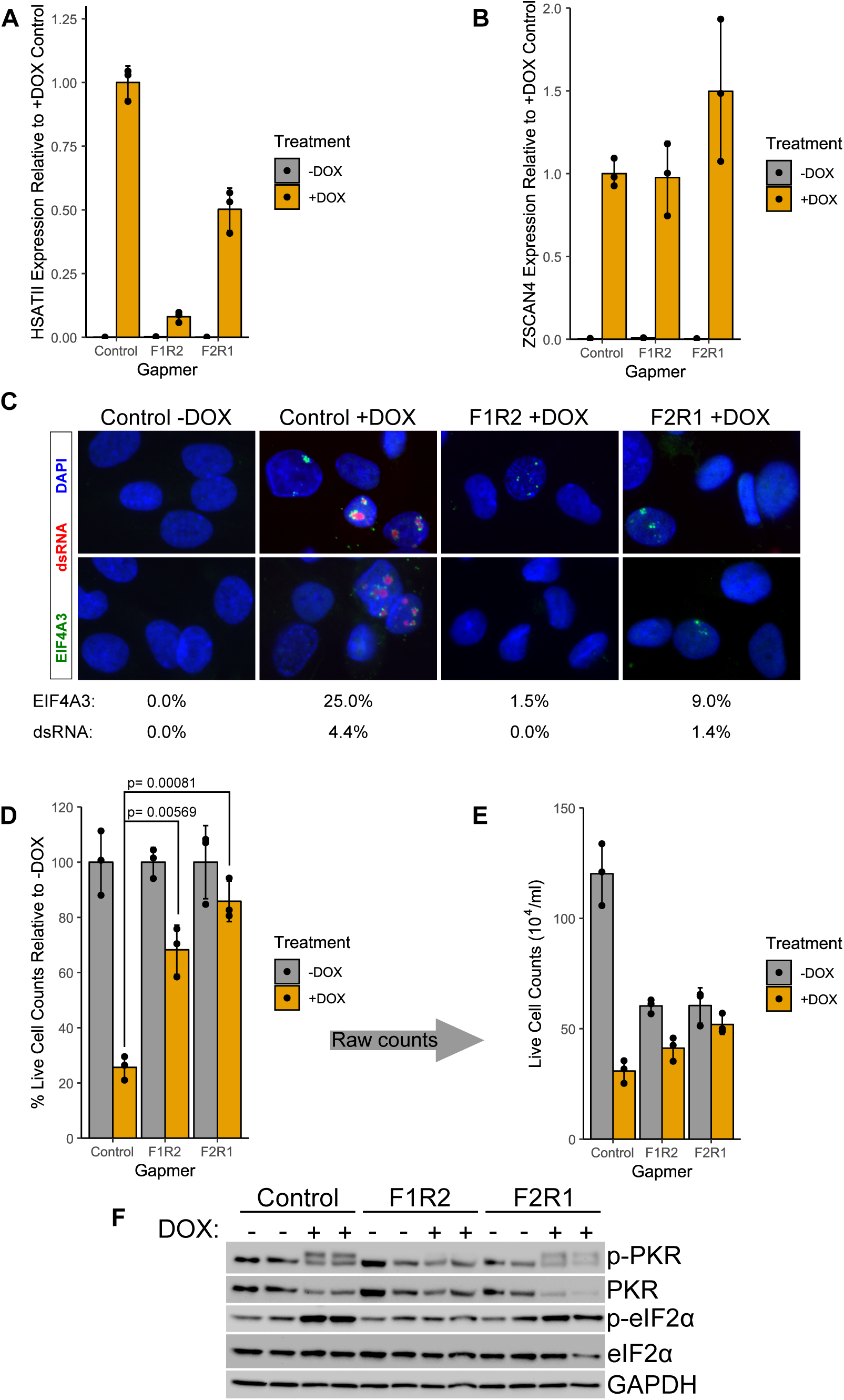
Gapmer-mediated knockdown of HSATII transcripts confirms their contribution to dsRNA foci and increases cell survival following DUX4 expression. (**A**) RT-qPCR showing levels of total HSATII RNA at 24 hours relative to the +DOX condition, normalized to RPL27A. Error bars represent the standard deviation of the mean for three independently cultured samples, shown as individual data points. (**B**) RT-qPCR showing levels of total ZSCAN4 (direct DUX4 transcriptional target) RNA at 24 hours relative to the +DOX condition, normalized to RPL27A. Error bars represent the standard deviation of the mean for three independently cultured samples, shown as individual data points (**C**) Representative examples of K1 (dsRNA) and EIF4A3 immunofluorescence images in MB135-iDUX4 cells at 18 hours -/+DOX. Percentages of K1 or EIF4A3 foci-positive nuclei in the indicated conditions are based on counts from > 500 randomly selected nuclei performed blinded to the experimental condition. Experiment was performed three times. (**D**) Trypan blue exclusion-based live cell counts in gapmer-transfected samples at 24 hours following DUX4 induction. The data are presented as the percentage of each gapmer’s paired -DOX sample for each time point. P-values were calculated using two-sided Welch’s T-test. No multiple-testing adjustment was used. Each dot represents the average relative live cell count from an independently cultured sample and error bars depict standard deviation from the mean. (**E**) Raw live cell counts used to make Fig. 5D which shows the reduction in live cell numbers due to HSATII knock-down (compare grey bars). Each dot represents the average live cell count from an independently cultured sample and error bars depict standard deviation from the mean. (**F**) Western blot showing levels of phosphorylated and unphosphorylated PKR and eIF2α in the indicated conditions as in Fig 5D. Each lane is an independently transfected sample. GAPDH serves as a sample loading control.

Our prior observation that activation of the dsRNA response contributed to DUX4-induced toxicity led us to hypothesize that HSATII transcription might be toxic to DUX4 expressing cells. Therefore, we next tested whether gapmer-mediated HSATII knockdown could decrease DUX4 toxicity. Indeed, as compared to a control gapmer, transfection of gapmers targeting HSATII forward and reverse transcripts enhanced survival following DUX4 expression when assessing the percentage of live cells compared to no DUX4 expression (Figure 5D), particularly with the F1R2 combination. HSATII gapmers also appeared to moderately retard myoblast proliferation leading to decreased numbers of total cells in the non-DUX4 expressing conditions (Figure 5E). This rescue of cell toxicity was similar to our previous report that knock-down of PKR or eIF2α partially rescued DUX4 toxicity (13). Importantly, gapmer transfection also diminished DUX4-induced, pro-apoptotic PKR phosphorylation (Figure 5F), most notably with the F1R2 gapmer combination, which occurs upon detection of dsRNAs by PKR. In addition, F1R2 and F2R1 gapmer combinations also diminished the increase of eIF2α phosphorylation that occurs in DUX4 expressing cells. As eIF2α is a phosphorylation target of activated PKR, this result confirms the reduction of PKR activity in DUX4 expressing cells following HSATII depletion. In sum, these data indicate that DUX4-induced HSATII transcripts are necessary for the observed intranuclear dsRNA foci formation and that depletion of HSATII transcripts decreases dsRNA-induced PKR phosphorylation and cell death following overexpression of DUX4.

### Endogenous DUX4 in FSHD muscle cells induces dsRNA-forming transcripts including HSATII

In cultures of FSHD skeletal muscle cells, *DUX4* is expressed in only a minor fraction of the muscle nuclei (1). To determine whether the expression of endogenous *DUX4* in FSHD muscle resulted in transcriptional upregulation of the identified dsRNA regions, we used RT-qPCR and found expression of genic as well as intergenic RNAs in the DUX4-induced dsRNA regions in both FSHD1 (MB073) and FSHD2 (MB200) cultured muscle cells, but not in control (MB135) muscle cells (Supplementary Material Figure S13). Sequencing cDNA amplicons from differentiated FSHD2 cells identified A-G mismatches compared to the gDNA, indicating ADAR editing and dsRNA formation of these transcripts in FSHD myotubes (Supplementary Material Figure S14).

We also found elevated HSATII expression in differentiated FSHD myotubes compared to a control non-FSHD cell line (Figure 6A). Consistent with our previous demonstration that a small proportion of FSHD muscle cells formed intranuclear dsRNA foci (13), RNA-FISH revealed rare intranuclear foci of HSATII using the probe to the reverse strand in FSHD myotubes, but not in control cells (Figure 6B). Therefore, as with ectopic expression of DUX4 in MB135-iDUX4 cells, endogenous DUX4 expression in FSHD myotubes also led to dsRNA expression and intranuclear foci of HSATII RNA.

**Figure 6.**
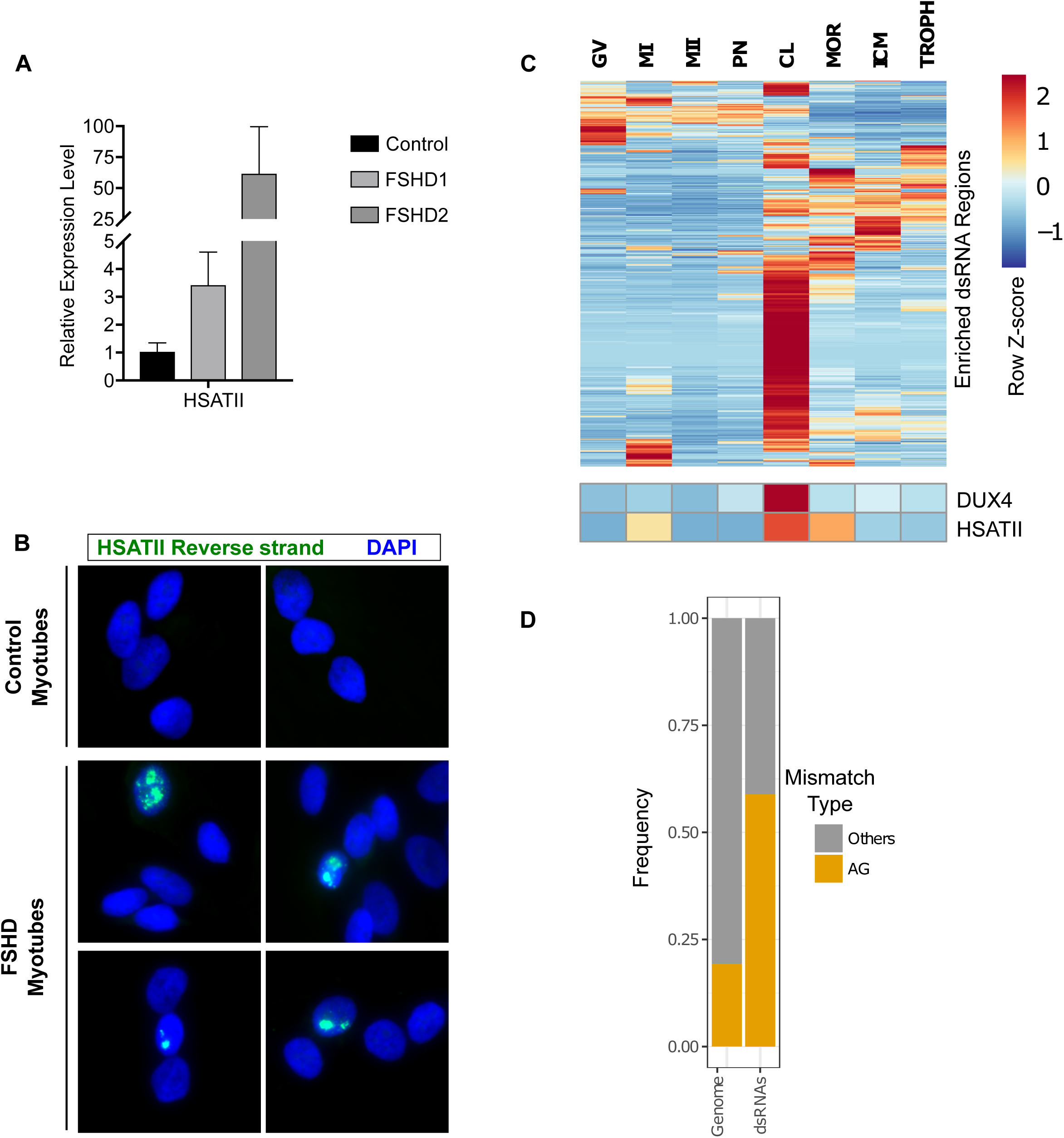
Endogenously expressed DUX4 induces dsRNA forming transcripts, including HSATII. (**A**) RT-qPCR data showing transcript expression levels of HSATII in control (MB135), FSHD1 (MB073) or FSHD2 (MB200) cells grown in differentiation medium. Data are normalized to RPL27 levels and shown relative to the control cell line. Data are depicted as the mean values of three experiments performed on independent days. Error bars represent the standard deviation of the mean. (**B**) RNA-FISH using probes targeting the reverse HSATII transcript in control (MB135) or FSHD (MB073) differentiated myotubes. Consistent with rare DUX4 expression in FSHD muscle cells, we observed a small subset of HSATII-positive nuclei in the FSHD cells, but not control cells. Images are representative from two independent experiments conducted on separate days. (**C**) Heatmap of DUX4-induced dsRNA, DUX4 and HSATII expression across various stages of human embryo development. Read counts were normalized and depicted as the within row Z-score. Data from Hendrickson, et al 2017 (3). GV (germinal vesicle), MI (metaphase I), MII (metaphase II), PN (pronuclear stage), CL (cleavage stage), MOR (morula) and ICM (inner cell mass), TROPH (trophectoderm). (**D**) Frequency plot of AG compared to all other possible mismatches in cleavage stage samples from Hendrickson, et al 2017 (3). Whole genome refers to all mismatches called, regardless of location and dsRNAs are mismatches called within DUX4-induced dsRNA regions.

### DUX4-induced dsRNA-forming RNAs including HSATII are expressed in pre-implantation embryos

Recent studies indicated that DUX4 and its mouse ortholog Dux activate a subset of the early cleavage stage transcription program in human and mouse embryos (2–4). In humans, EGA involves the expression of HSATII and HERVL retrotransposons and in mice involves the expression of MERVL and GSAT major pericentric satellite repeats. To determine whether cleavage stage expression of *DUX4* correlated with dsRNA formation, we re-analyzed published early embryo RNA-seq data (3) across several different stages of development. As previously reported, we identified *DUX4* expression at the cleavage stage of embryonic development and this correlated with increased expression of many of the same DUX4-induced dsRNA-forming regions in myoblasts, including HSATII (Figure 6C and Table S4). Analysis of base mismatches between the RNA-seq and the reference genome sequence revealed a dramatic enrichment of A-G mismatches over other mismatch types across DUX4-induced dsRNA regions (Figure 6D). Interestingly, both forward and reverse strand transcripts of HSATII were at near equal levels in the cleavage stage (Supplementary Material Figure S15), perhaps because the samples contained a heterogeneous pool of embryos at different cleavage cell numbers. Thus, our analysis verified that many of the DUX-induced dsRNA-containing transcripts identified in DUX4 expressing myoblasts, including HSATII, are also present in the early embryo.

## Discussion

When DUX4 is expressed in induced pluripotent stem cells or skeletal muscle cells, it activates genes characteristic of a totipotent developmental expression program (2–4). For example, DUX4 activates the expression of *ZSCAN4* which has roles both in telomere elongation (37) and DNA demethylation through the degradation of UHRF1 and DNMT1 (38), as well as *KDM4E*, which is a lysine demethylase that can relieve chromatin repression (39). DUX4 also activates the expression of repetitive elements, including LINE-1, HERVL and MaLR endogenous retroviruses, and HSATII satellite repeats (3, 4, 8, 9, 28). In this study, we show that DUX4 creates intergenically derived dsRNAs from a subset of these repeats. Most dramatically, bidirectional transcription of HSATII repeats results in the formation of intranuclear dsRNA foci and gapmer knockown of the HSATII RNAs partially rescues DUX4 toxicity together with decreasing the DUX4-induced phosphorylation of PKR and eIF2α. These results demonstrate that the toxic dsRNA response induced by DUX4 is mostly mediated by the induction of HSATII transcripts. Although we used forced expression of DUX4 in skeletal muscle cells to identify these dsRNA-forming transcripts, analysis of RNA and RNA-seq showed that these dsRNA-forming regions were also transcribed and ADAR-edited in FSHD muscle cells and cleavage stage embryos expressing the endogenous DUX4. Along with our previous results (13), these findings solidify a mechanism of dsRNA formation in FSHD that can contribute to cell toxicity.

In mice, during the cleavage stage of early embryogenesis, pericentric “major satellite” repeats are expressed first in the forward, then reverse direction (relative to the consensus sequence) where they have been hypothesized to form dsRNAs that direct repressive heterochromatin modifications to the pericentromeres (36, 40). However, the mechanism of regulating these transcripts remained unknown. Gapmer-mediated depletion of these repeat transcripts blocked chromocenter formation and interrupted development at the 4-cell stage, indicating a necessary role for satellite RNAs in establishing pericentric heterochromatin (36). While transcription of satellite repeats appears to be necessary for heterochromatin establishment, whether dsRNA is required is less clear (41). As with DUX4 in human embryogenesis, mouse Dux is expressed in the cleavage stage and, interestingly, is coincident with transcription of GSAT (major satellite) pericentric repeats (3, 4). In the case of HSATII, our study shows that DUX4 appears to first initiate HSATII transcription in predominantly one direction, producing an RNA encoding the reverse complement of the consensus HSATII sequence, and then later induces HSATII transcription in the opposite direction coincident with the formation of multiple distinct intranuclear dsRNA foci. Given that this bidirectional expression pattern of HSATII by DUX4 parallels the temporal dynamics of mouse expression of GSAT repeats, we postulate that DUX4, and by analogy mouse Dux, has a primary role in regulating the bidirectional transcription of pericentric satellite repeats in early human and mouse embryogenesis.

In contrast to its potential function in the early embryo, the expression of HSATII RNA in FSHD muscle could have both direct and indirect biological consequences that could alter RNA processing and chromatin repression. Previously we reported that DUX4 expression induces intranuclear focal aggregates of EIF4A3 (13), and here we showed that EIF4A3 accumulates with nuclear HSATII RNA foci. Although it remains unclear whether the association of EIF4A3 with HSATII RNA is due to splicing of the HSATII transcripts or another recruitment mechanism, the formation of EIF4A3 foci correlates with the inhibition of nonsense mediated RNA decay (NMD). This suggests that the sequestration of EIF4A3 might contribute to DUX4-mediated inhibition of NMD, and could act in concert with the DUX4-mediated depletion of UPF1 (13, 42). Similar to the association of EIF4A3 with the predominantly expressed reverse HSATII strand, the association of ADAR1 foci with HSATII dsRNA could also contribute to toxicity in DUX4-expressing cells. For example, ADAR1 aggregation might limit the enzyme’s activity on non-HSATII containing endogenous dsRNAs, which would be expected to lead to activation of cellular antiviral pathways. In accordance with this, *ADAR1* knockout leads to dsRNA-associated apoptosis in a human cell line that can be rescued by simultaneous knockout of *RNASEL* (43), paralleling our previous finding that DUX4 toxicity can be alleviated by knockdown of *RNASEL* (13).

DUX4-mediated induction of HSATII RNA might have other consequences as well. HSATII transcription has been associated with the formation of intranuclear foci of components of the PRC1 complex (polycomb bodies), which can potentially lead to de-repression of LINE-1 elements or other repeats (30). The DUX4-induced formation of polycomb bodies, as indicated by the BMI1 foci, are consistent with a model where HSATII transcription in DUX4 expressing cells might have a similar role in the formation of polycomb bodies and activation of LINE-1 repeats. The co-localization of the BMI1 foci with phospho-H2AX foci suggests that the DNA breaks might be the driver of the location of the BMI1 foci, although further work will be necessary to determine whether the DNA damage is a consequence of HSATII repeat expression. In this regard it is interesting to note that, in mice, expression of pericentric major satellite repeats can lead to genome instability, likely by destabilizing DNA replication forks (35). This causes activation of the DNA damage response and cell death which can be alleviated by knockout of *p53*. Whether HSATII expression causes these effects in human cells is currently unknown, but its demonstrated activation in tumor cell lines (33, 44) suggests that its expression may similarly trigger genome instability.

A possible function for non-HSATII intergenic dsRNAs induced by DUX4 remains more speculative. It is interesting to note the relative enrichment of LINE-1 and ERV repeats within many of the intergenic DUX4-induced dsRNAs, classes of repeats that are expressed in the cleavage stage embryo, but generally repressed in most tissues (45). Similar to the proposed role of HSATII repeats, the double-stranded secondary structure of these transcripts might also facilitate heterochromatin formation. This has been postulated as a function of dsRNA-forming lncRNAs, which can recruit methyltransferase activity to DNA (46). Interestingly as well, the DUX4-induced long intergenic transcripts often contain a region of dsRNA enrichment and are reminiscent of the previously characterized “very long intergenic non-coding” (vlinc) RNAs observed in a multitude of pluripotent cells and also cancer lines (47, 48), though the nature of the transcriptional regulation of these vlincRNAs is presently not well understood.

The enrichment of retroelement-derived dsRNAs in a subset of DUX4 target genes might also serve a function. Although retroelements are typically subject to DNA methylation and other silencing mechanisms (49), their presence can have a positive effect on gene expression at early stages of development (50–52). Silencing of these retroelements at later stages might then provide precise temporal control of DUX4 targets whose continued expression would counteract later differentiation processes. It is possible, therefore, that the dsRNAs in DUX4-induced genes might contribute to the subsequent silencing of these genes through regional dsRNA-mediated heterochromatin formation.

In summary, our work confirms the induction of dsRNA containing transcripts by DUX4. Similar to constitutively expressed dsRNAs, many of the DUX4-induced dsRNAs are enriched for inverted *Alu* repeats in genic transcripts. In contrast to constitutive dsRNAs, a large fraction of DUX4-induced dsRNAs are intergenic and are modestly enriched for LTR-containing ERVs and LINE-1 elements, but highly enriched for HSATII satellite repeats. DUX4 induction of HSATII repeats in a bidirectional, temporally controlled pattern mirrors the dynamics of mouse major satellite repeat expression in the early embryo and suggests that human DUX4 activation of HSATII repeat transcripts in the cleavage stage embryo might play a similar role in establishing pericentric heterochromatin via dsRNA formation. Our results from depleting DUX4-induced HSATII transcripts indicate that HSATII dsRNAs contribute to DUX4 toxicity in somatic cells and might also contribute to DUX4 toxicity in FSHD and is consistent with our prior observation that knockdown of RNASEL or PKR also partially rescues DUX4-toxicity (13). The expression of DUX4 and bidirectional HSATII transcripts in the early embryo suggests a role for these dsRNAS in early development that merits further study.

## Materials and Methods

### Cell culture

De-identified human primary myoblast cell lines from the Fields Center for FSHD Neuromuscular Research at the University of Rochester Medical Center were immortalized by retroviral transduction of CDK4 and hTERT (53). All immortalized human myoblasts were cultured in F10 medium (Gibco/ThermoFisher Scientific) supplemented with 20% FBS (GE Healthcare Life Sciences) and 1% penicillin/streptomycin (Gibco/ThermoFisher Scientific), 10.0 ng/ml recombinant human FGF (Promega) and 1.0 μM dexamethasone (Sigma-Aldrich). Myoblasts were differentiated by culturing in DMEM (Gibco/ThermoFisher Scientific) containing 1% horse serum (Gibco/ThermoFisher Scientific), 1% penicillin/streptomycin (Gibco/ThermoFisher Scientific), 10 μg/ml insulin and 10 μg/ml transferrin for 48-72 hours. All “+DOX” labels mean that cells were grown for approximately 18 hours in the presence of 1.0 μg/ml of doxycycline hyclate (Sigma-Aldrich), unless otherwise noted.

### Gapmer LNA transfections

Gapmers were transfected into MB135-iDUX4 cells on the day prior to doxycycline induction using Lipofectamine RNAiMAX reagent (ThermoFisher Scientific) following the manufacturer’s instructions. Gapmers were ordered from QIAGEN. Sequences of gapmers are shown below (a ‘+’ indicates a locked nucleic acid modification in the following base):

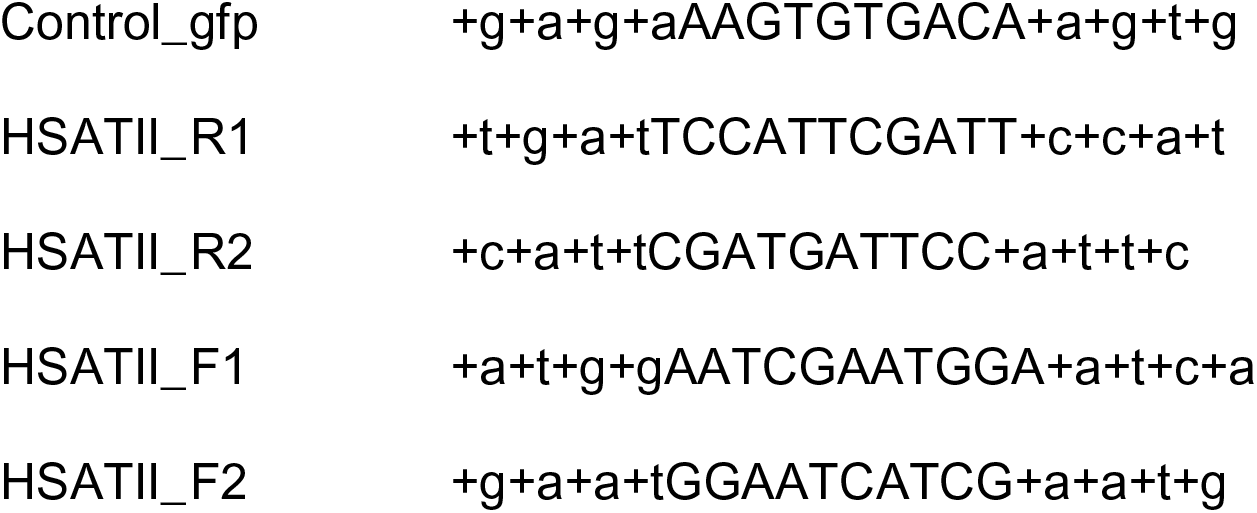

|  |  |  |
| --- | --- | --- |
| 499 | Control_gfp | +g+a+g+aAAGTGTGACA+a+g+t+g |
| 500 | HSATII_R1 | +t+g+a+tTCCATTCGATT+c+c+a+t |
| 501 | HSATII_R2 | +c+a+t+tCGATGATTCC+a+t+t+c |
| 502 | HSATII_F1 | +a+t+g+gAATCGAATGGA+a+t+c+a |
| 503 | HSATII_F2 | +g+a+a+tGGAATCATCG+a+a+t+g |

### dsRNA Immunoprecipitation

Human myoblast MB135-iDUX4 cells were treated -/+DOX and washed in PBS, trypsinized and counted prior to lysis. Approximately 1.2×10^6^ cells were used for each IP. Lysis was performed in 1.0 ml total volume by sonication using a Diagenode Bioruptor on light setting (5 minutes total, 30s on/off at 4°C) in a buffer composed of 15 mM Tris pH 7.5, 0.1 M NaCl, 5 mM MgCl_2_, 0.5% Triton X-100, 1 mM dithiothreitol, and 40 U/ml RNase inhibitor (ThermoFisher Scientific). Lysates were precleared using 40.0 μl of protein G Dynabeads (ThermoFisher Scientific) for 1 hour prior to an overnight incubation at 4°C with either J2, K1 or an isotype-matched anti-GFP (IgG) control antibody. 40.0 μl of protein G Dynabeads were added the following morning for 1 hour to bind the antibody, and beads were subsequently washed four times with 1.0 ml of cold lysis buffer. After the final wash, 1.0 ml of TRIzol (ThermoFisher Scientific) was added directly to the beads for RNA extraction.

### RNA extraction and library preparation

TRIzol RNA extractions were performed as per manufacturer’s recommendations. Ribosomal RNA depletion was performed using the NEBNext rRNA depletion kit (New England Biolabs). First strand synthesis was achieved using SuperScript III reverse transcriptase (ThermoFisher Scientific) with random hexamers and using the following thermocycler conditions: 25°C 10min, 50°C 30min, 55°C 30min and 85°C 5min. Second strand synthesis was achieved using the NEB second strand cDNA synthesis kit, following manufacturer’s recommendations. The double-stranded cDNA was fragmented to approximately 250 bp average size using a Diagenode Bioruptor sonicator in 120 μl total volume for three 10-minute intervals on medium intensity at 4°C, 20s on/off, replenishing with pre-cooled water between each interval. Libraries were made from the AmpureXP (Beckman Coulter) purified cDNA using the Ovation Ultralow system V2 kit and following the manufacturer’s instructions. Stranded total RNA-seq libraries were made from 1.0 μg of MB135-iDUX4 RNA using the KAPA stranded RNA-seq kit with RiboErase (HMR) and following the manufacturer’s instructions. Libraries were sequenced using 100 bp single-end sequencing on the Illumina HiSeq 2500 platform by the FHCRC Genomics facility.

### dsRIP sequence alignment and differential peak calling

We essentially treated our dsRIP-seq data as diffuse (e.g. histone mark) ChIP-seq to find locations of enrichment. First, dsRNA-IP reads were aligned to hg38 using BWA version 0.7.12 with the option -n 6. Next, peaks were called by comparison to the IgG background control condition using MACS2 version 2.1.0 using the options: --broad, --broad-cutoff 0.01, -q 0.01. We then used Diffbind version 2.3.8 to differentially call dsRNA enrichment between +/-DOX conditions with “minOverlap” set at 0.5, which treated J2 and K1 antibodies as replicates for comparison purposes. The significance threshold was set at 1e-10. We extracted the consensus peakset from the Diffbind dba report, setting the significance threshold as 1 to capture all dsRNA-enriched loci. We filtered our list of enriched dsRNAs to eliminate dsRNAs which overlapped known rRNAs, tRNAs and snRNAs (obtained from UCSC table browser repeatMasker track filter repclass: “snRNA OR rRNA OR tRNA OR snoRNA”) using bedtools “intersect”. We used the Integrative Genomics Viewer (IGV) version 2.3.98 for data visualization. We used the “kpPlotRegions” function from the karyoploteR package to plot location of DUX4-induced dsRNA-enriched regions.

### Feature annotation

We used bedtools “annotate” and GenCode v24 feature annotations downloaded from the UCSC genome browser to classify per-base feature overlaps of dsRNAs with “intergenic” being defined as the absence of a feature overlap and “downstream” and “upstream” being defined as 5 kb 3-prime or 5-prime of an annotated gene feature, respectively. Overlapping features were prioritized as: coding exons > 5 UTR > 3 UTR > intron > upstream 5 kb > downstream 5 kb > intergenic.

### Stranded RNA-seq

Stranded RNA-seq libraries were aligned using TopHat version 2.1.1 with Bowtie version 2.2.9 and the iGenomes UCSC gene transfer format file with the following options: --read-mismatches 8 --read-edit-dist 8.

### RNA editing detection

We used the python script, REDItools *denovo* version 1.0.4 (54) to call editing sites on our aligned samples. Replicate BAM files were merged using samtools before analysis. The following REDItools parameters were used to limit false positives: -E (Exclude positions with multiple changes), -a t (two-tailed Fisher’s exact test), -c 10 (minimum read coverage of 10), -T 6-6 (6 bases were trimmed from each end of the reads), -q 25 (minimum quality score of 25), -n 0.1 (minimum editing frequency of 0.1) and -v 3 (minimum number of reads supporting the variation set at 3). The SNP147 GTF file was downloaded from the UCSC genome table browser to exclude known polymorphic sites from consideration. We used a p-value threshold value of < 0.1.

### Repeat analysis

For repeat counting in our stranded, ribosome-depleted RNA-seq dataset we used the TEtranscripts script from TEtoolkit package (55) with the multi-mode option and a custom GTF file which was made from the current (07-Nov-2016) release of Dfam non-redundant repeat matches. This approach enabled counting within predicted HSATII repeats (which are not available in the hg38 repeatMasker annotation used). We performed differential expression analysis on the resulting count table with DESeq2 using default options. For differential expression of repeats in our dsRIP-Seq data, we performed the above analysis in K1 or J2 versus IgG in doxycycline-treated samples.

### Confirmation of A-I editing

We amplified dsRNA regions containing putative A-I RNA editing sites using Phusion High Fidelity DNA polymerase (ThermoFisher Scientific) from genomic DNA or cDNA of doxycycline-induced MB135-iDUX4 cells. Purified amplicons were Sanger sequenced by the FHCRC genomics facility on a 3730xl DNA Analyzer and visualized using Geneious Pro software, version 5.0.4.

### RNA isolation and real time qPCR

For immunoprecipitation samples, RNA was isolated using TRIzol reagent (ThermoFisher Scientific) according to the manufacturer’s protocol and using GlycoBlue (ThermoFisher Scientific) as a co-precipitant. For all other samples, RNA was isolated with the RNeasy kit (QIAGEN), according to the manufacturer’s protocol. Purified RNA was treated with DNaseI (ThermoFisher Scientific), heat inactivated, and reverse transcribed into cDNA using Superscript III (ThermoFisher Scientific) following the manufacturer’s instructions. Quantitative PCR was performed with SYBR green reagent (ThermoFisher Scientific). For RT-qPCR of RNA immunoprecipitation, random hexamers (ThermoFisher Scientific) were used for non-strand specific detection. Data were calculated relative to +DOX input and normalized to U2 background RNA values for each independent immunoprecipitation. For HSATII forward strand specific RT-qPCR, we used the ‘SATII_pericentric_1L’ oligo to prime cDNA synthesis with a 5-prime linker ATGGATCACGAGAACACTGA sequence. For qPCR we used the ‘SATII_pericentric_1R’ and linker sequence as the primers. Samples were subsequently normalized to U2 signal in the respective random hexamer primed reactions, as above.

### HSATII RNA-FISH and combined immunofluorescence RNA-FISH

Locked nucleic acid, FITC conjugated HSATII probes were purchased from QIAGEN and are based on the sequence used in previous publications (30, 56, 57). Probe 1: 5’ FAM-ATTCCATTCAGATTCCATTCGATC detects the reverse HSATII transcript. Probe 2, which detects the forward HSATII transcript is the reverse complement of probe 1 with 5’ FAM modification. LNA positions were not disclosed. We performed HSATII RNA-FISH essentially as described previously (30) with modifications. Briefly, cells grown on coverslips were rinsed in cold CSK buffer (100 mM NaCl, 300 mM Sucrose, 3mM MgCl_2_, 10mM Pipes pH 6.8) and then permeabilized in ice cold extraction buffer (CSK, 0.1% TX-100 and 10 mM Ribonucleoside Vanadyl Complex (NEB)) for 5 minutes. After a second CSK rinse, cells were fixed in 4% paraformaldehyde (Electron Microscopy Sciences) in PBS for 10 minutes at room temperature. Cells were dehydrated in series of 70% ethanol, 90% ethanol and 100% ethanol and then air dried. Probes were diluted to 5.0 pmol/ml and denatured at 75°C for 5 minutes in WCP buffer (50% formamide, 2XSSC, 10% dextran sulfate) and hybridized overnight at 37°C. Coverslips were washed with 50% formamide/2xSSC at 37°C and then twice with 2xSSC/0.1% TX100 for 5 minutes before mounting using prolong gold antifade with DAPI (ThermoFisher Scientific).

For combined immunofluorescence and RNA-FISH we first fixed cells on coverslips using 4% paraformaldehyde for 7 minutes at room temperature. Cells were permeabilized with 0.5% TX100 in PBS for 5 minutes prior to overnight incubation with primary antibodies at 4°C. Samples were incubated with appropriate secondary antibody for 1 hour at room temperature prior to fixation of the antibody interaction via a second crosslinking step using 4% paraformaldehyde for 10 minutes at room temperature. Cells were incubated with probes which were denatured as above for 2 hours at 37°C prior to sequential washes with 15% formamide 2xSSC for 20 minutes at 37°C, 2xSSC for 20 minutes at 37°C and 2xSSC for 5 minutes at room temperature. Cells were imaged using a Leica TCS SP5 II confocal microscope, where indicated, or a Zeiss AxioPhot. Image channel merging and processing was performed using ImageJ software.

### Analysis of human embryo RNA-seq data

We aligned raw reads to genome build hg38 using TopHat2, allowing two mismatches per read and maximal twenty multiple alignments. The counts for enriched dsRNA regions were computed by summarizeOverlaps() from the GenomicAlignments Bioconductor package with “IntersectStrict” mode. Next, we applied the countRepeats R package (https://github.com/TapscottLab/countRepeats), which is designed specifically for counting reads of repetitive elements originating from forward/reverse transcripts. To obtain DUX4 counts, we aligned the raw reads to the customized D4Z4 “genome” comprised of DUX4 exon sequences.

### Cloning of HSATII and testing DUX4 activation

a ∼1.5 kb PCR product was cloned from MB135 cell genomic DNA using the “SATII pericentric” primers which were synthesized with flanking NheI and HindIII restriction sites such that the amplicon could be readily ligated into the pGL3 basic vector. The opposite orientation clone was made using the original vector as template by PCR with the flanking restriction sites swapped to reverse the orientation of the HSATII insert in the same position. We tested for luciferase expression by RT-qPCR using SYBR green reagent and the comparative C_T_ method.

### Antibodies (product ID, epitope, company)

J2 anti-dsRNA, SCICONS English & Scientific Consulting; K1 anti-dsRNA, SCICONS English & Scientific Consulting; ab180573 anti-EIF4A3, Abcam; anti-hADAR, polyclonal rabbit antibody obtained from B. Bass; ab126783 anti-BMI1, Abcam; JBW301 anti-phospho-histone H2A.X (S139), Millipore; D7F7 anti-PKR, Cell Signaling Technologies; ab32036 anti-phospho-PKR (T446), Abcam; sc-133132 anti-eIF2α, Santa Cruz Biotechnology; ab32157 anti-phospho-eIF2α (S51), Abcam; GTX28245 anti-GAPDH, GeneTex.

### Primers

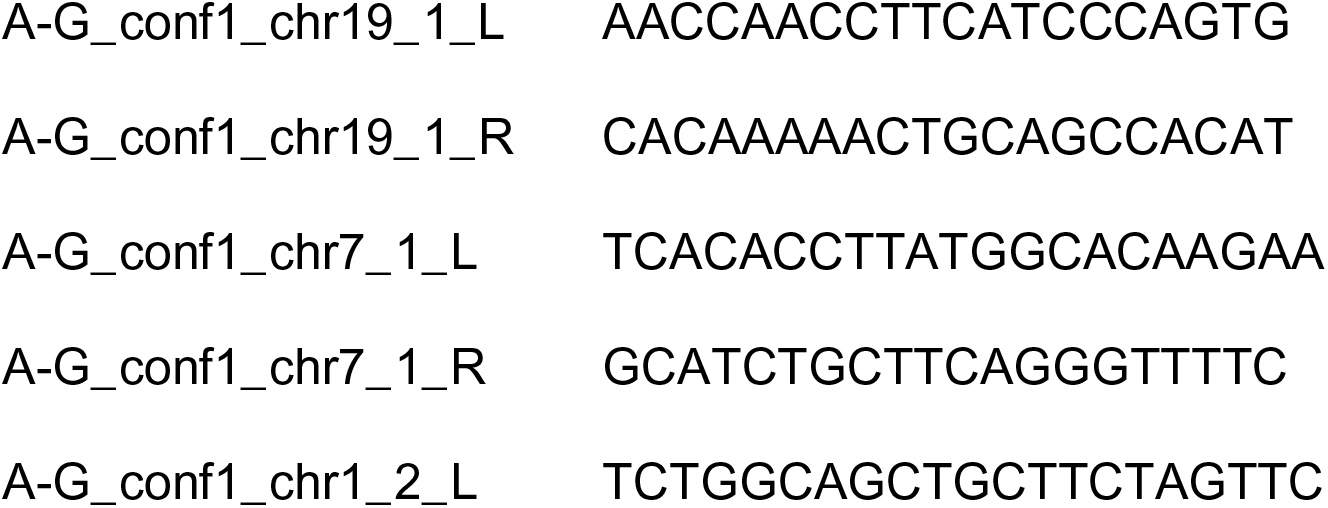
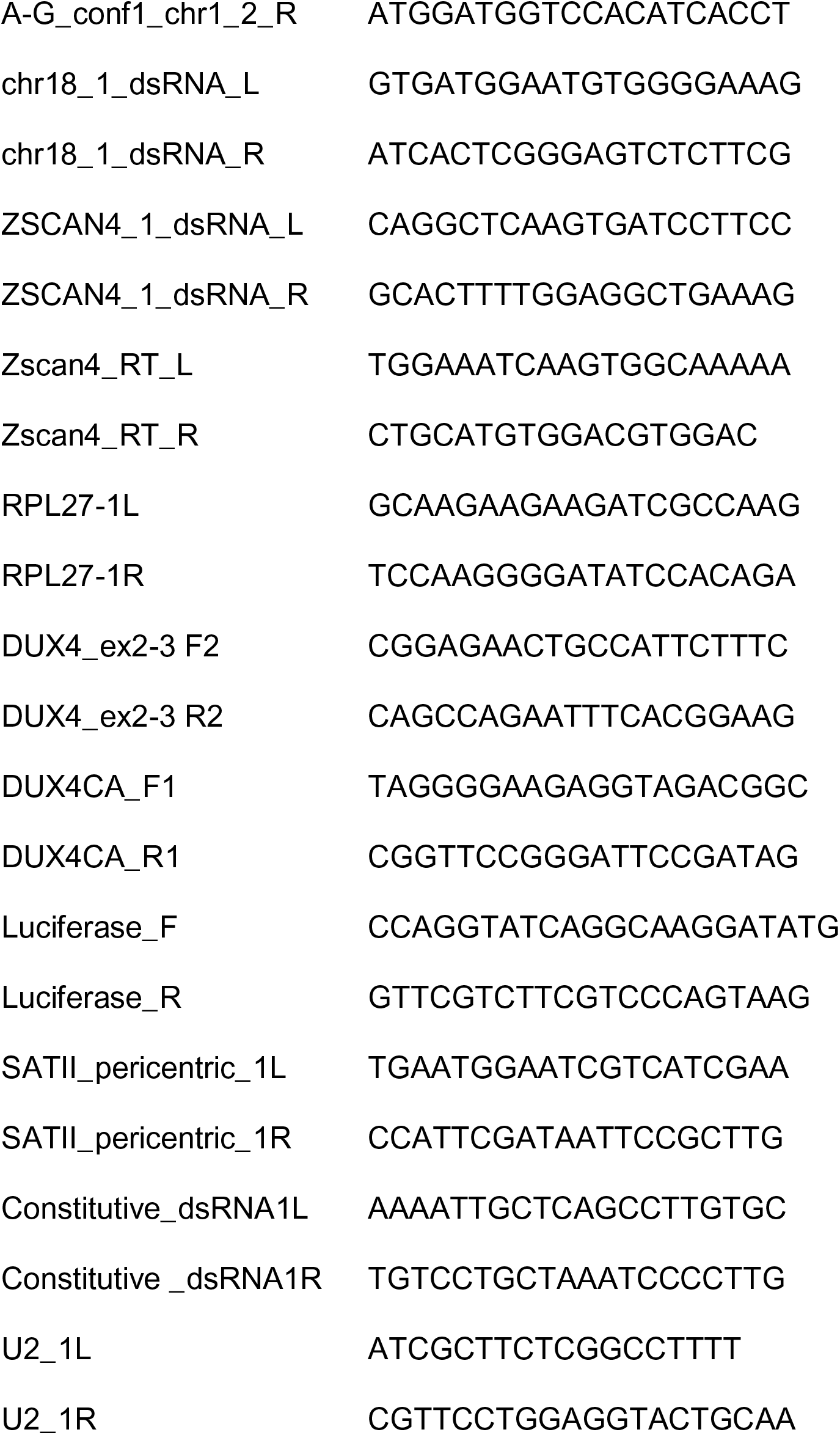

## Data Availabilty

MB135-iDUX4 dsRIP-seq and total stranded RNA-seq data were deposited into the Gene Expression Omnibus (GEO) under the SuperSeries accession number GSE114940. DUX4 ChIP-seq data used in this study are available from GEO under accession number GSE33838. Early embryo RNA-seq data were obtained from GEO accession number GSE72379.

## Acknowledgements

We thank Mathew Thayer for helpful comments and assistance with the FISH protocol and Lisa Kursel for assistance with confocal imaging. We thank Rebecca Resnick, James Thomas and Christine Beck for critical reading of the manuscript and Bob Eisenman for helpful comments.

## Supporting Information Legends

**Table S1. Differential expression analysis of regions enriched for dsRNA.** Table showing filtered DiffBind differential enrichment results of K1 and J2 versus IgG MACS2 derived peaks in +DOX compared to -DOX condition in MB135-iDUX4 cells.

**Table S2. Differential expression of repeats in DUX4 expressing MB135 myoblasts.** DESeq2 differential expression results from ribosome depleted total RNA-seq of repeat counts in the +DOX compared to -DOX conditions in MB135-iDUX4 cells.

**Table S3. dsRNA enriched repeats.** DESeq2 differential expression results of repeat counts in the +DOX condition of MB135-iDUX4 dsRIP-seq with K1 or J2 RIPs compared to IgG RIP. Table is divided into four sheets showing: 1. DUX4-induced repeats that are enriched by K1 versus IgG, 2. all K1 enriched repeats, 3. DUX4-induced repeats that are enriched by J2 versus IgG, and 4. all J2 enriched repeats.

**Table S4. Normalized read counts and FPKM of DUX4, HSATII repeats and DUX4-induced dsRNAs in human embryo developmental RNA-seq dataset.** Read counts and FPKM values used to create the heatmap in Figure 6C.

