## Supplemental Figures for "DUX4-induced bidirectional HSATII satellite repeat transcripts form intranuclear double stranded RNA foci in human cell models of FSHD"

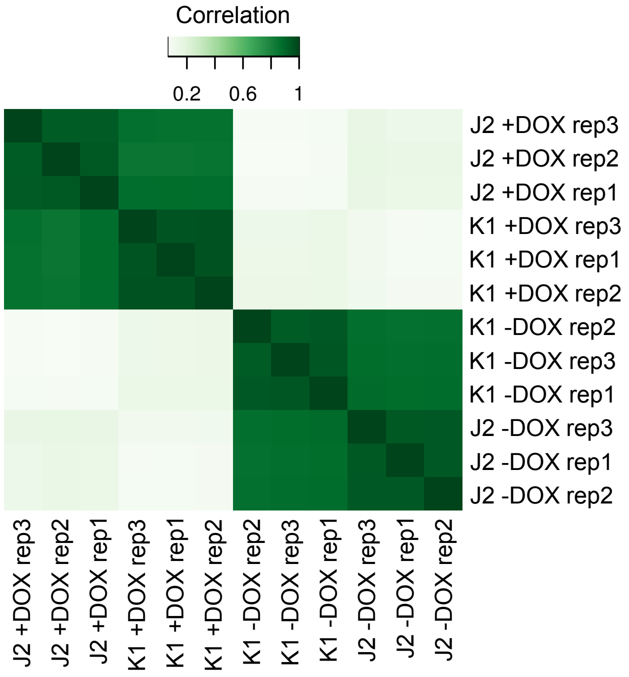


**Figure S1. Correlation heatmap based on affinity scores across all consensus peaks.** Generated using the DiffBind Bioconductor package. There is a high correlation between K1 and J2 antibody binding within each condition, which formed the basis for merging the two separate datasets in subsequent analyses.

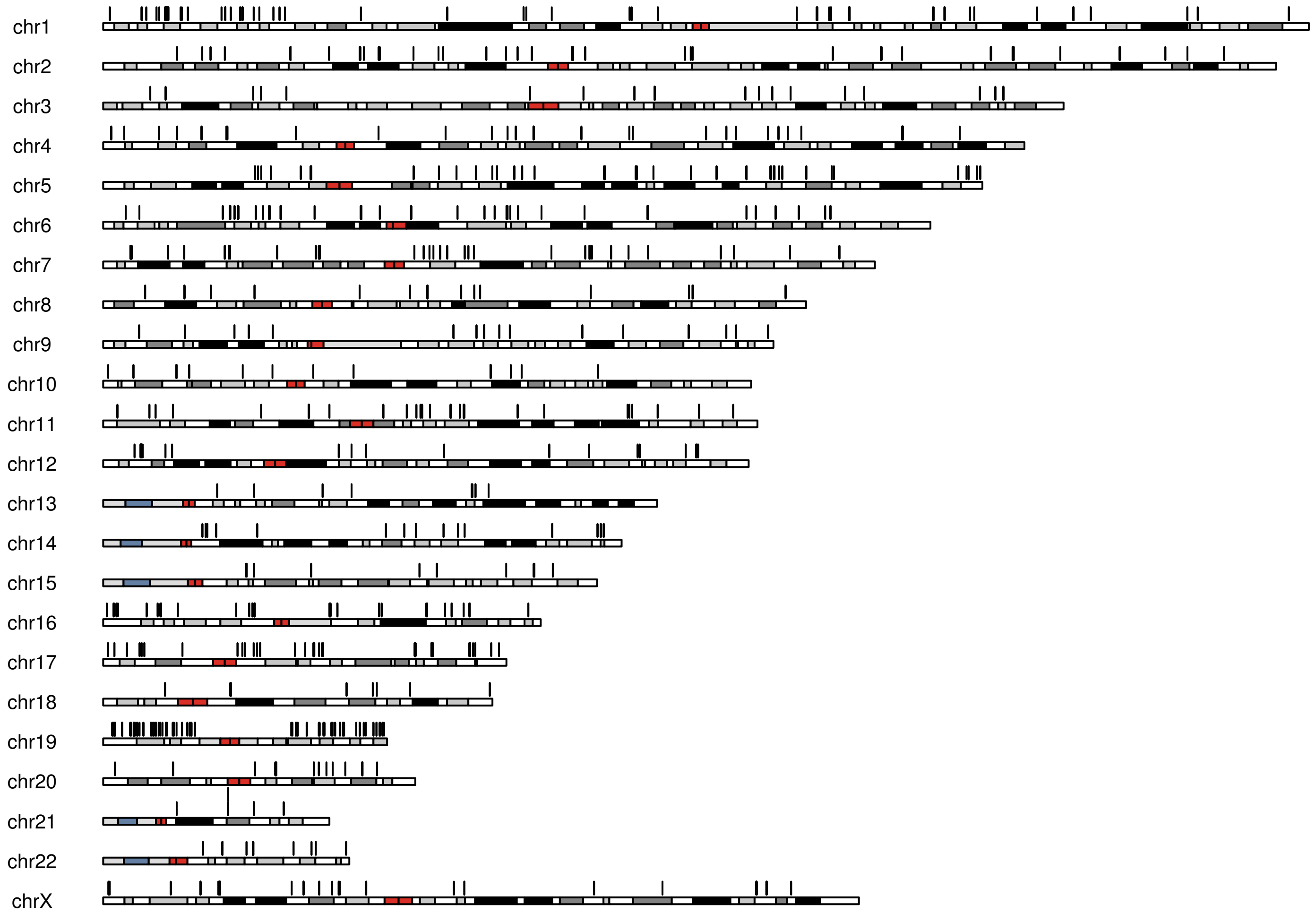


**Figure S2 . Chromosomal ideogram of DUX4-induced dsRNA locations.**
Representation of all DUX4-induced dsRNA locations in MB135-iDUX4 cells (black bars above chromosomes).

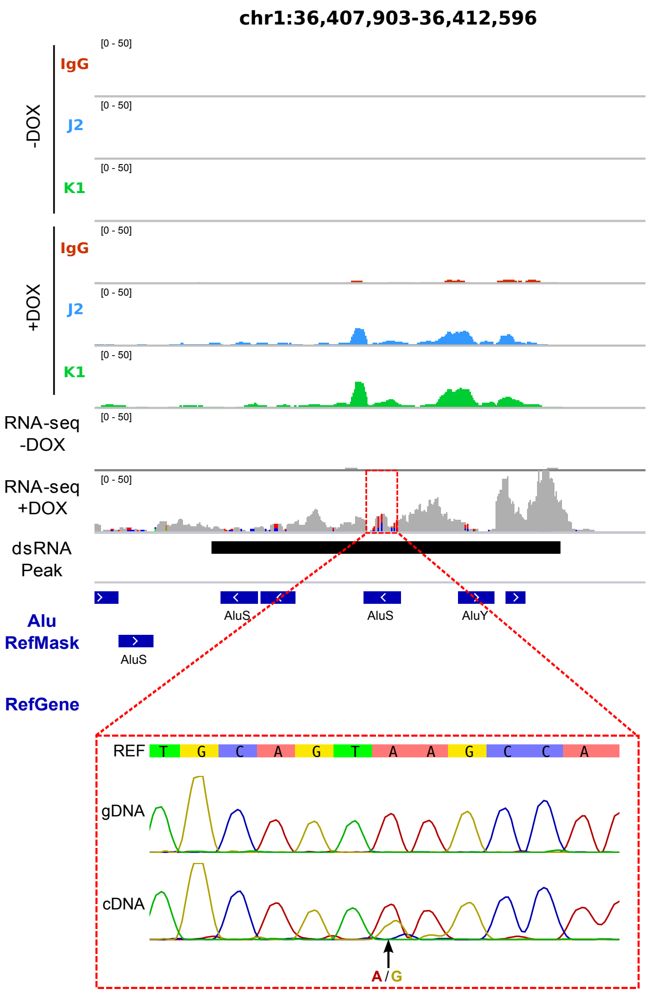


**Figure S3. Sanger sequencing confirmation of an additional RNA-editing event in a DUX4-induced dsRNA.**

See Fig 1D-1E for description.

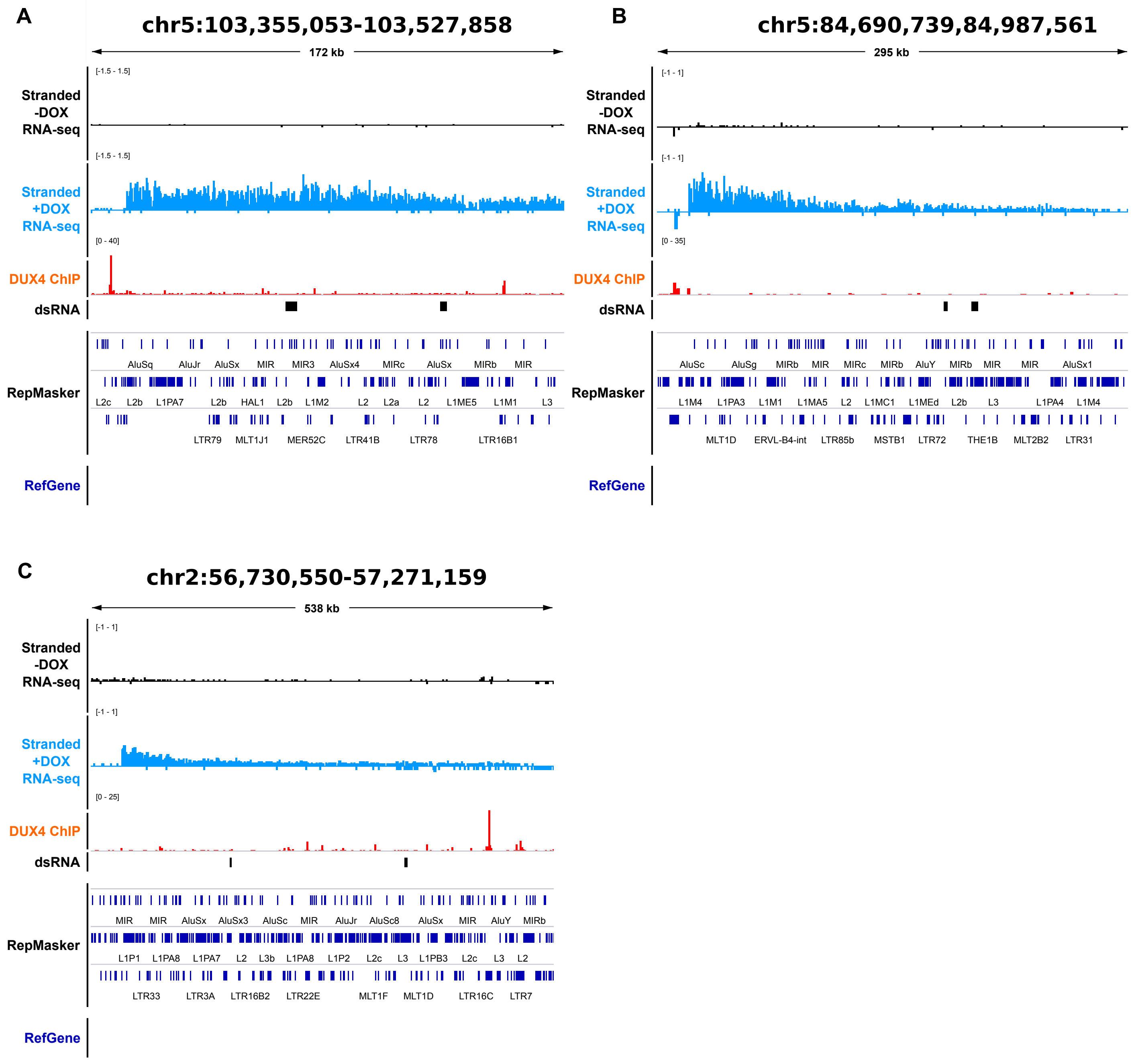


**Figure S4. Additional examples of DUX4-induced intergenic dsRNAs.**
(A-C) As in Figure 2C-2D, for the indicated locations.

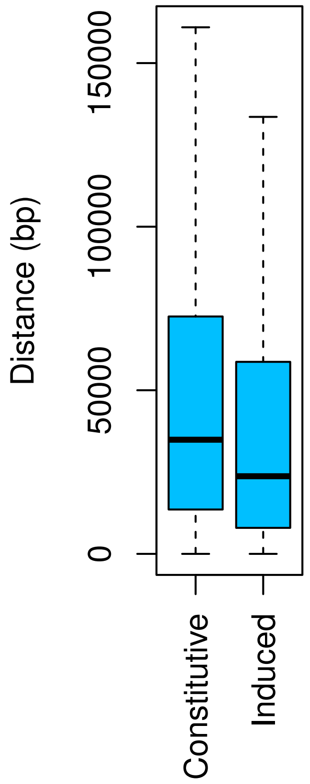


**Figure S5. DUX4-induced dsRNAs are more proximal to DUX4 ChIP-seq peaks than constitutive dsRNAs.**
Boxplot of base pair distance to the nearest DUX4 ChIP-seq peak of the indicated dsRNA types. P = 6.83e-10 based on the Wilcoxon rank sum test. For plotting purposes only, extreme outliers (points beyond whiskers) were omitted.

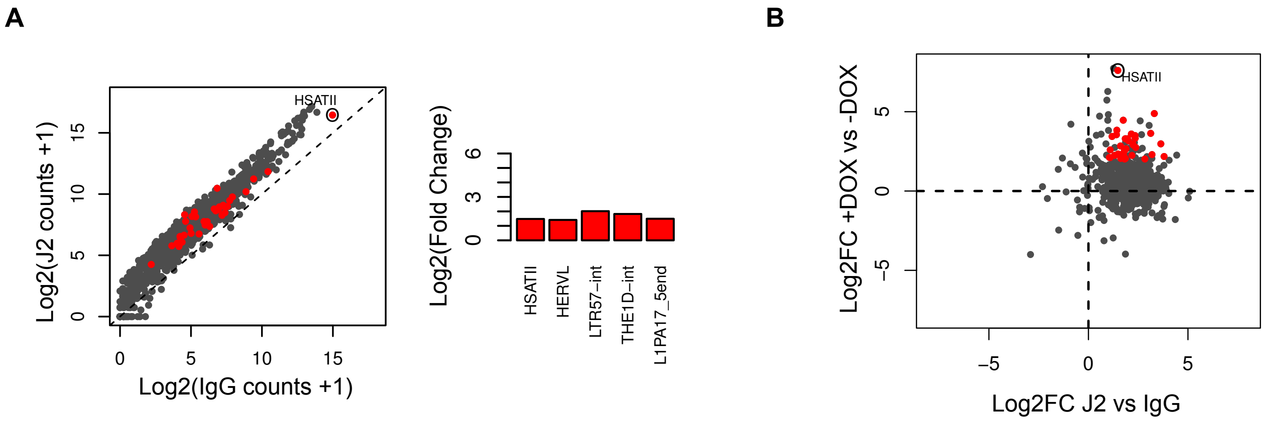


**Figure S6. DUX4-induced HSATII repeat enrichment in J2 immunoprecipitations.**
(A) As in Fig 3B except using J2 antibody data. (B) As in Fig 3C except comparing to J2 antibody moderated log2 fold change values.

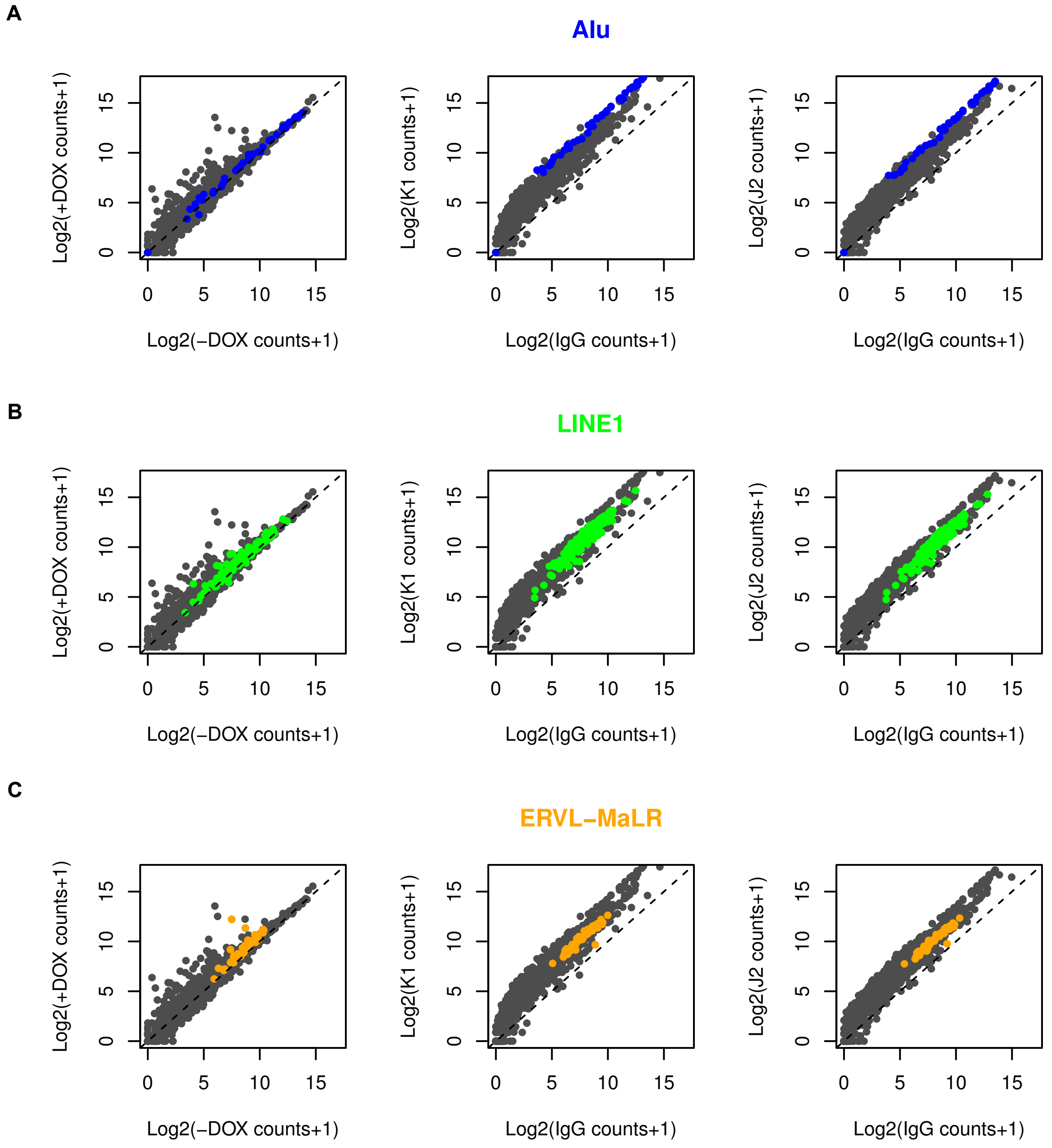


**Figure S7. dsRNA enrichment of DUX4 activated repeat families.**
(A) Scatterplot depicting log2 normalized read counts of Dfam predicted subfamilies within stranded RNA-seq dataset of MB135-iDUX4 cells -/+ DOX (left), +DOX K1 vs IgG dsRIP samples or +DOX J2 vs IgG samples, with Alu repeats highlighted in blue (without expression or significance thresholding). (B) As in (A) except highlighting LINE1 repeat subfamilies in green. (C) As in (A) except highlighting ERV-MaLR repeat subfamilies in orange.

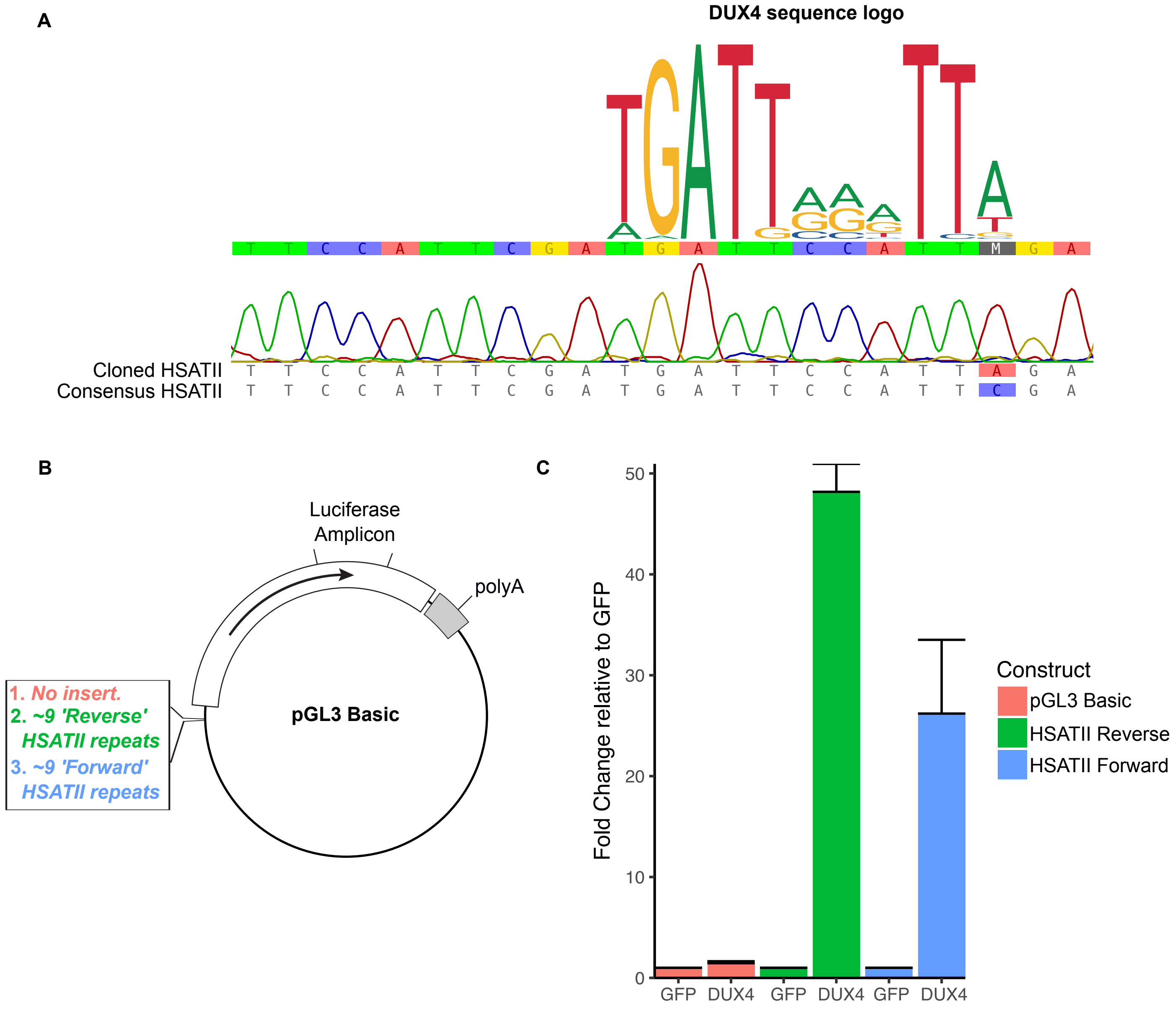


**Figure S8. DUX4 directly activates HSATII transcription.** (A) Example Sanger sequencing from cloned ~1.5 kb HSATII amplicon showing divergence from consensus HSATII sequence that creates a good match for DUX4 binding. Sequence logo is based on the JASPAR database DUX4 motif. (B) Schematic showing location of ~1.5 kb HSATII cloning site in pGL3 Basic vector, upstream of luciferase gene and amplicon site for primers used to detect luciferase expression. (C) RT-qPCR showing levels of luciferase expression normalized to RPL27 and depicted relative to GFP transfection control for the indicated pGL3 Basic luciferase construct. HEK293T cells were transfected with either GFP or DUX4 expression vectors and co-transfected with pGL3 Basic (no insert), HSATII Reverse or HSATII Forward vector. Here "forward" means that the HSATII amplicon cloned into the pGL3 Basic vector is in the consensus orientation relative to the luciferase sense strand. Error bars represent the standard deviation of the mean for three independent transfections except in the case of the pGL3-basic DUX4 transfection where one sample was omitted due to a technical failure which led to non-amplification.

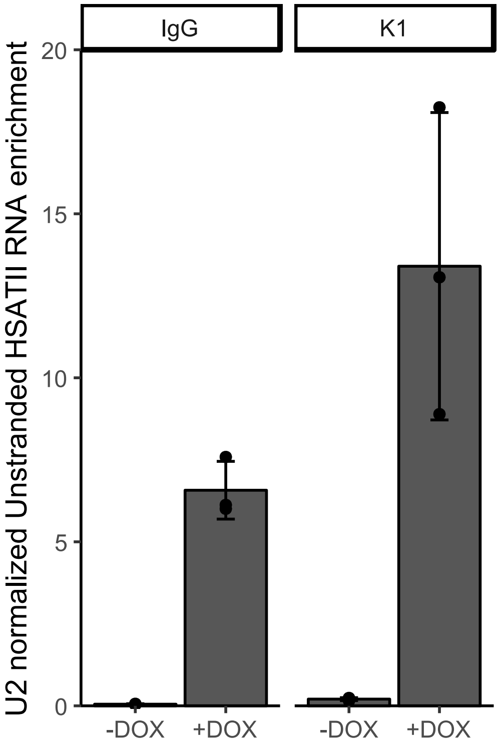


**Figure S9. Non-strand specific RT-qPCR for HSATII following dsRIP.**
As in Fig 4E except a non-strand specific RT-qPCR was performed to detect all (forward and reverse) HSATII transcripts.

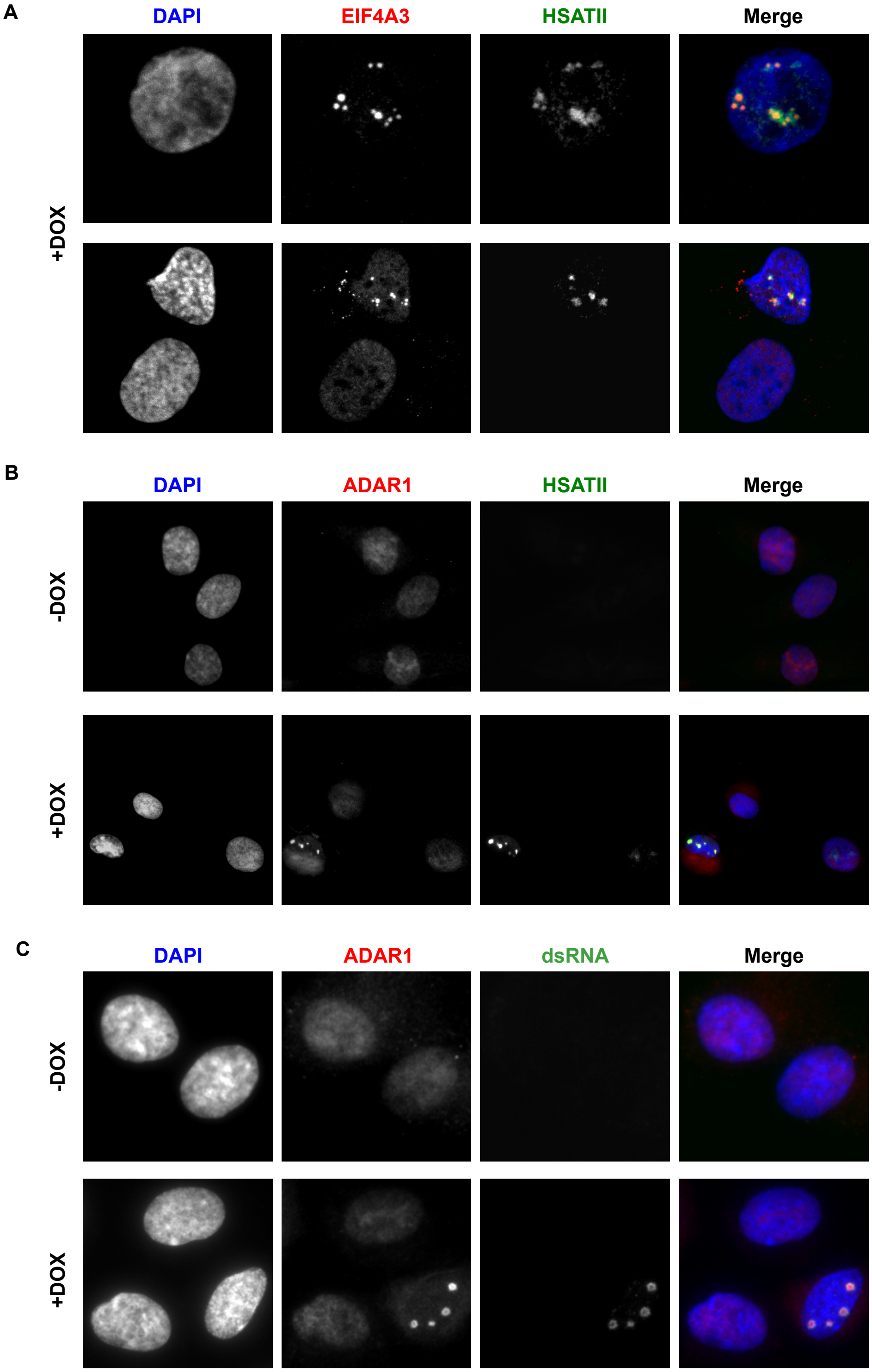


**Figure S10. RNA binding proteins aggregate with the HSATII containing dsRNA foci in DUX4 expressing cells.**

(A) Immunofluorescence in MB135-iDUX4 cells using antibody for EIF4A3 combined with RNA-FISH using probes targeting the reverse HSATII transcript. All cells were induced with doxycycline for approximately 18 hours. Images are representative from two independent, combined IF RNA-FISH experiments conducted on separate days. (B) Combined immunofluorescence with ADAR1 antibody and RNA-FISH with probes that detect the reverse HSATII strand transcripts in MB135-iDUX4 cells -/+ DOX. Images are representative from one experiment. (C) Co-immunofluorescence with K1 anti dsRNA and ADAR1 antibodies in MB135-iDUX4 cells -/+ DOX. Images are representative from three independent co-IF experiments conducted on separate days.

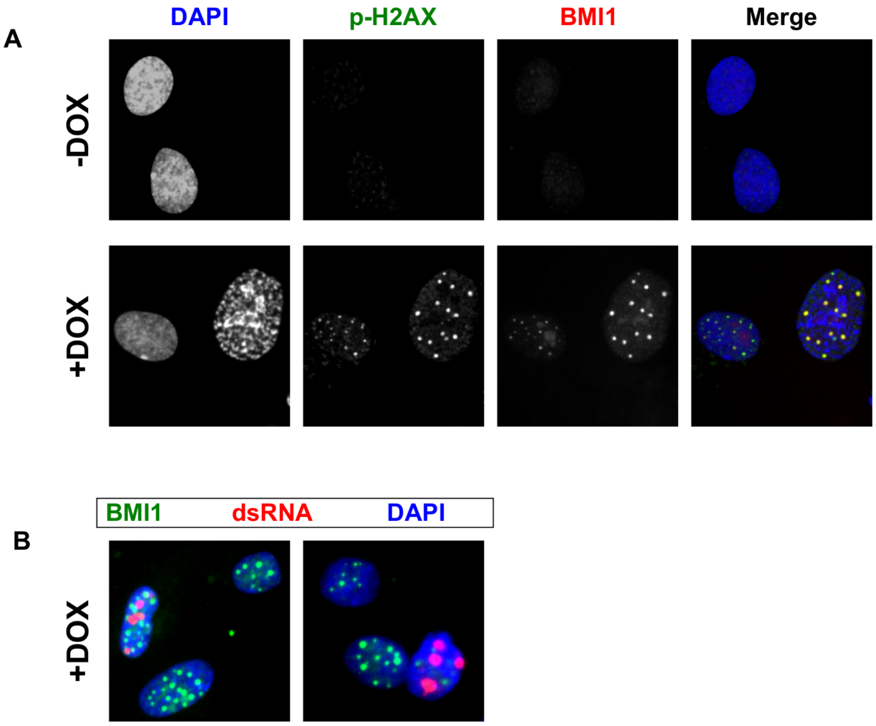


**Figure S11. BMI1 aggregates in DUX4 expressing cells do not colocalize with DUX4-induced dsRNA aggregates.**
(A) BMI1 and phospho-H2AX co-immunofluorescence in -/+ DOX treated MB135-iDUX4 cells. Images are representative from three independent experiments conducted on separate days. (B) BMI1 and K1 dsRNA antibody co-immunofluorescence in doxycycline treated MB135-iDUX4 cells. Images are representative from one experiment.

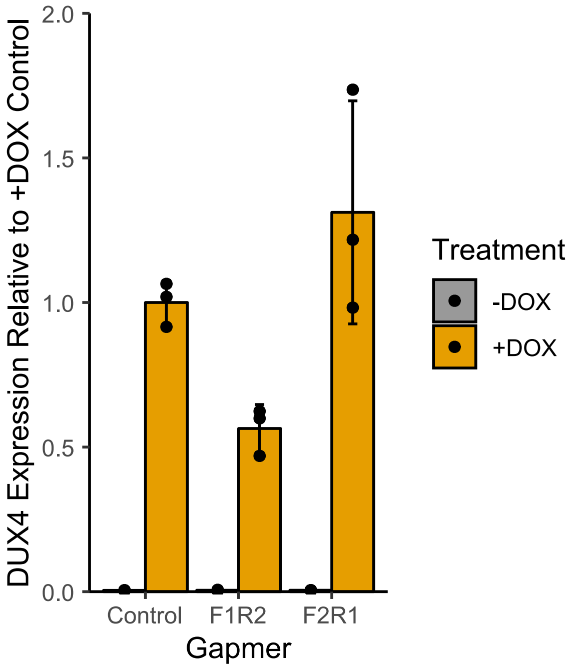


**Figure S12. Related to Figure 5A-B.**
RT-qPCR showing levels of DUX4 transgene RNA relative to the +DOX condition, normalized to RPL27A. Error bars represent the standard deviation of the mean for three independently cultured samples, shown as individual data points.

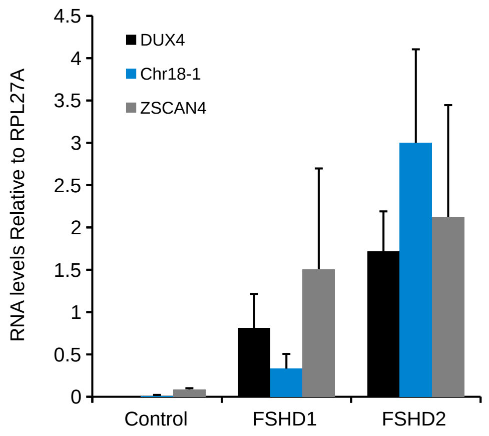


**Figure S13. RT-qPCR of additional identified DUX4-induced dsRNA enriched locations.**
RT-qPCR data showing transcript expression levels using indicated primers in control (MB135), FSHD1 (MB073) or FSHD2 (MB200) cells grown in differentiation medium. Data are normalized to RPL27 levels and depicted as the mean values of three experiments performed on independent days. Error bars represent the standard deviation of the mean.

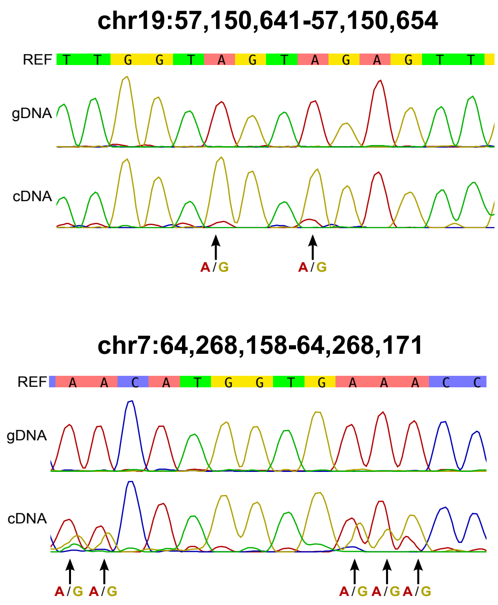


**Figure S14. Confirmation of dsRNA editing in FSHD cells.**
Sanger sequencing results of amplicons designed across the shown regions (hg38) in either genomic DNA (gDNA) or complementary DNA (cDNA) of differentiated FSHD2 (MB200) cells, with AG mismatches highlighted.

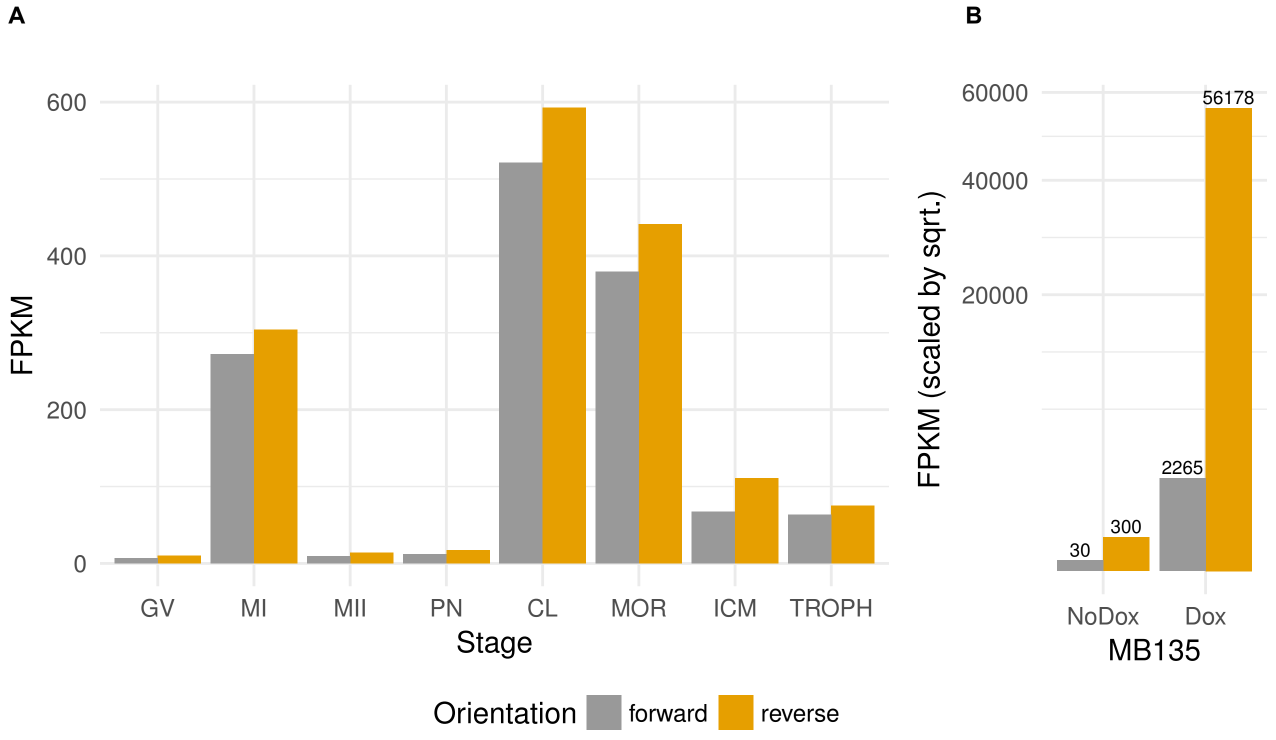


**Figure S15. HSATII transcript strand expression.**
(A) FPKM from RNA-seq data in the indicated developmental stages of reads mapping to HSATII repeat sequences on either forward or reverse strand (relative to the consensus HSATII sequence). (B) as in (A) except in MB135-iDUX4 cells and scaled by the square root, with unadjusted FPKM values depicted above bars.
